# Sustainable cattle management by communities supports African wildlife

**DOI:** 10.1101/2025.10.09.681397

**Authors:** Erin Connolly, Holly A.I. Pringle, Omiros Pantazis, Guilherme Braga Ferreira, Emily K. Madsen, Daniel J. Ingram, Taras Bains, Gabriel J. Brostow, Sarah Carroll, Georgia Cronshaw, Paul De Ornellas, Enrico Di Minin, Robert M. Ewers, Kevin Gichangi, Oisin Mac Aodha, Martin Mulama, Muthoni Njuguna, Liam Pattullo, Alastair Pickering, Alexandre Rabeau, Marcus Rowcliffe, Fiona Spooner, Liam Thomas, Yussuf Wato, Emily Woodhouse, Ben Collen, Georgina M. Mace, Kate E. Jones

**Author notes:** Co-first authorship (Erin Connolly, Holly A.I. Pringle). Deceased (May 2018) (Ben Collen). Deceased (September 2020) (Georgina M. Mace).

## Abstract

Community-based conservation (CBC) initiatives aim to reconcile biodiversity protection with local livelihoods, yet their effectiveness in protecting wildlife remains uncertain, often hinging on local management^1,2^. We evaluated a globally significant CBC model in Kenya’s Greater Maasai Mara Ecosystem (GME), where conservancies, run jointly by Maasai landowners and the tourism sector, employ rotational cattle grazing to support both wildlife and pastoralism^3,4^. Using a ∼1200 km^2^ grid of 180 camera traps across gradients of livestock pressure in Maasai Mara National Reserve and three conservancies in 2018, we collected and analysed over 2 million images with a customised AI-powered pipeline. We found a positive impact of observed cattle pressure on mammal community occupancy and species richness, except for at the highest levels of cattle grazing. However, sheep and goat grazing and proximity to infrastructure had a negative impact. These results provide evidence that wildlife and pastoralism can coexist under community-led stewardship^5^, but only with active management and targeted control of emerging threats. AI tools such as our image classifier may contribute to more adaptive community-led management of these areas^6^. As conservation policy shifts beyond formal protected areas, our findings support CBC as a scalable model for conserving biodiversity within working landscapes, offering a pathway to meet global targets while maintaining local livelihoods^7^.

---

Community-based conservation (CBC) is increasingly promoted as a cornerstone of meeting global biodiversity goals, including protecting 30% of the Earth by 2030^7,8^. Aiming to align biodiversity conservation with human wellbeing, CBC devolves management rights to local communities and allows multiple land uses, while strict protected areas (PA) typically restrict resource access^1^. Though strict PAs remain essential to safeguard some of the world’s most threatened species, CBC is often positioned as a more equitable and scalable alternative which supports local livelihoods and rights^1^. Yet despite growing policy attention, whether CBC sustains biodiversity at scale is poorly understood, particularly in Africa, where two-thirds of conserved land lies outside strict PAs and the average size of vertebrate populations has declined by 76% in the last 50 years^9^. In shared African rangelands, CBC must reconcile wildlife conservation with livestock production, which is both essential to pastoralism and a leading driver of biodiversity loss^10^. Research has explored primarily social outcomes of CBC^1,2^, while ecological evidence remains limited and often species- or trophic-group-specific^11,12^, potentially obscuring how entire wildlife communities respond to multiple anthropogenic pressures across working landscapes.

Kenya’s Greater Maasai Mara Ecosystem (GME) offers a compelling case study for evaluating the effectiveness of CBC in conserving wildlife across a dynamic landscape. This savanna contains one of the richest assemblages of wild megafauna globally and marks the northernmost point of the Serengeti-Mara wildebeest and zebra migration^13^. Conservation areas in the GME are unfenced. The Maasai Mara National Reserve (MMNR) is strictly protected and no livestock grazing is permitted, though illegal night grazing occurs^14^. MMNR is surrounded by community-based conservancies, which were found to hold up to 83% of all GME wildlife^15,16^. Conservancies operate through partnerships between Maasai landowners and tourism operators, whereby landowners receive lease payments in exchange for minimising settlement on these land parcels and limiting cattle grazing to rotational management plans^4^. Sheep and goats (shoats), however, are not actively managed in conservancies; shoats are permitted to graze near settlements but may also graze illegally outside of these areas. Savannas such as the GME currently face mounting pressures from human sedentarisation, infrastructure development, and livestock expansion^17–20^. Thus, the GME provides a unique opportunity to disentangle the impacts of multiple livestock species and other anthropogenic pressures, offering critical insights into the ecological effectiveness of CBC.

Here, we quantify how three anthropogenic pressures: cattle grazing, shoat grazing, and distance to human infrastructure, shape occupancy and species richness of 35 wild mammal species across approximately 1200 km^2^ of the GME (Figure 1, Extended Data Figure 1; Methods). This study spans gradients of these pressures across four conservation areas: a portion of the strictly protected MMNR known as Mara Triangle (510 km^2^), and three surrounding community-based conservancies: Mara North (320 km^2^), Naboisho (220 km^2^), and Olare-Motorogi (150 km^2^). We deployed 180 camera traps from 9 October to 25 November 2018 and used a customised deep learning algorithm, the Maasai Mara Classifier (MMC), to classify our camera trap dataset of over 2 million images (Figure 2, Supplementary Table 1, Extended Data Figure 2; Methods). We implemented hierarchical multispecies occupancy models (MSOMs) and species richness measures to assess how these pressures affect a diverse assemblage of mammals (Figure 1; Methods). Our analyses provide insights into how wildlife communities respond to co-occurring anthropogenic pressures, highlighting the ecological outcomes of CBC efforts across a dynamic socio-ecological system.

**Figure 1:**
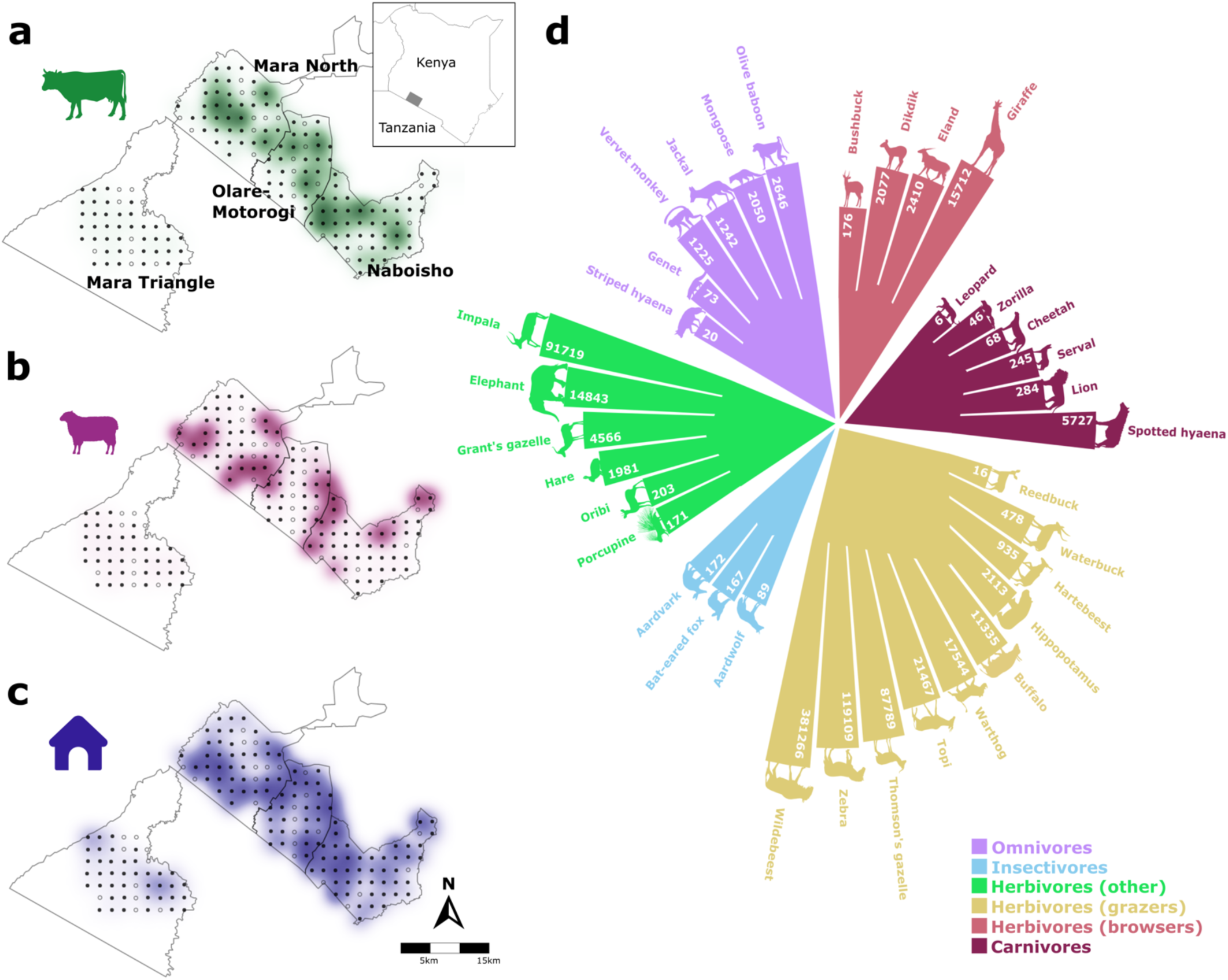
Biodiversity monitoring across anthropogenic pressure gradients in the Greater Mara Ecosystem, Kenya. A-C: Spatial distribution of three anthropogenic pressures. Black dots represent locations of 147 camera traps (CTs) used in our analyses; empty dots represent 33 camera traps excluded due to malfunction or theft. Darker colours denote higher pressure (Extended Data Figure 1). A: Cattle trap rate [(number of detection events x 100)/number of full days the camera was operational] (green) at each CT station (range: 0-157.14). B: Shoat (sheep and goat combined) trap rate (purple) at each CT station (range: 0-194.44). C: Distance to human infrastructure (blue) from each camera trap station (range: 94-10,574 m). D: Circular bar plot of 35 wild mammal species detected by camera traps and classified using a deep learning model (Methods). Species are grouped by dietary functional classes (Extended Data Table 1). Numbers inside bars indicate total observations per species; bar length representing log-transformed observations. Functional groups are colour coded to match bars. Silhouettes obtained from PhyloPic (www.phylopic.com).

**Figure 2:**
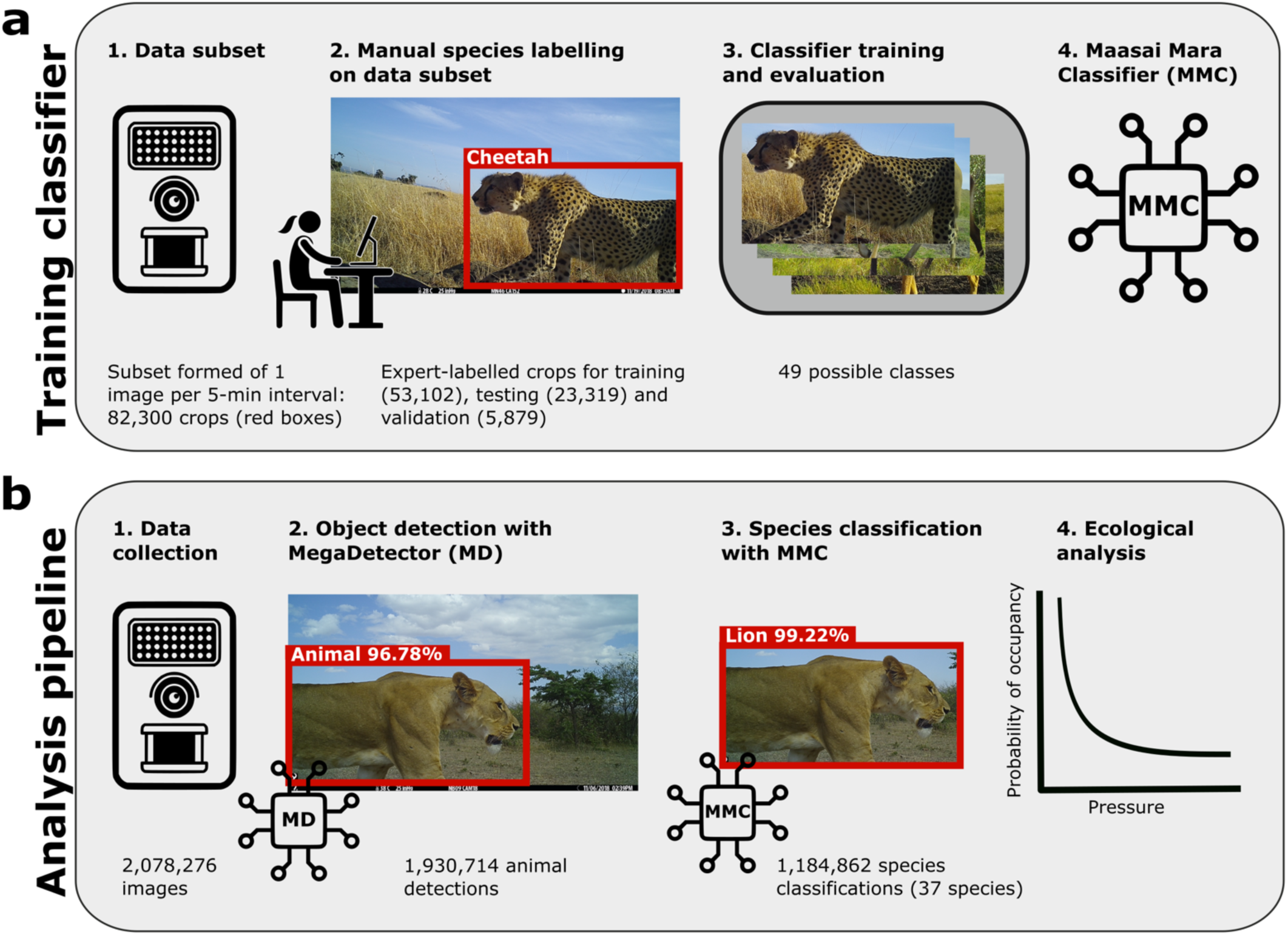
Camera trap data processing workflow. A: Training the Maasai Mara Classifier (MMC). Experts labelled a subset of images to create training, testing, and validation sets (∼9:4:1 ratio). B: Data analysis pipeline. To classify our full dataset, we applied MegaDetector (version 4)^71^ (MD), extracting animal crops. We then applied the MMC, which classified crops to species level. Finally, after confidence thresholding, we fed the species predictions from the MMC into our occupancy model to investigate mammal responses to anthropogenic pressure gradients.

## Managed cattle can coexist with wildlife

Our multispecies occupancy model provides strong posterior support for a positive association between mammal community occupancy and cattle grazing (posterior mean = 0.36; 95% Credible Interval (CrI) = 0.06, 0.65, Figure 3A). We also found strong posterior support for a negative quadratic (nonlinear) effect (mean = -0.09; 95% CrI = -0.17, -0.004) (Figure 3B, Extended Data Table 1). While both have strong support, the linear effect size is much larger, and explains marginally more of the community occupancy response (linear f = 0.99, quadratic f = 0.98), indicating that conservancies are broadly effective in fostering cattle-wildlife coexistence at the levels of pressure we captured. The smaller negative quadratic effect suggests that there are localized areas, at 92% of observed pressure and above, where cattle pressure becomes high enough to reduce wildlife occupancy (Figure 1A, Extended Data Table 2). Together, these responses suggest that conservancies are successfully supporting both cattle and wildlife populations under management, but further increases in cattle pressure could risk the sustainability of this coexistence. Estimated species richness followed a similar pattern (Figure 3C), indicating that at the time of study, cattle grazing intensities broadly support coexistence, but elevated pressure may erode mammal diversity.

**Figure 3:**
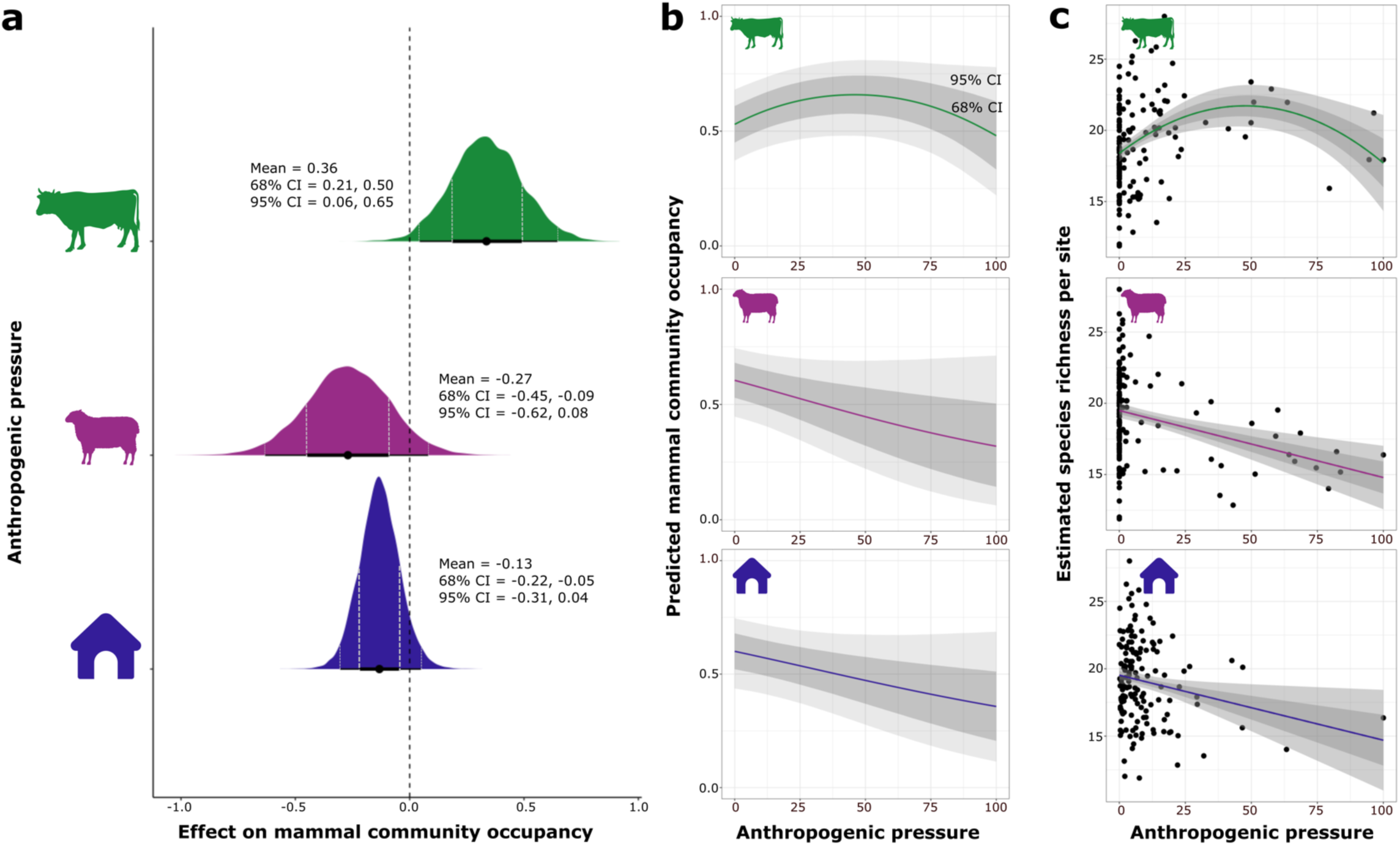
Responses and predictions of mammal community occupancy along three anthropogenic pressure gradients. A: Coefficient plot showing relative effect sizes of three standardised anthropogenic pressures on mean mammal community occupancy. Points represent effect size, thin horizontal bars with vertical dashed lines represent 95% credible intervals (CrIs) and thick horizontal bars with vertical dashed lines represent 68% CrIs. Effects are considered significant at a given level if that CrI does not overlap zero. The y-axis of each distribution corresponds to the density of posterior draws at that particular effect size value. B: Means (coloured lines) and CrIs of predicted mammal community occupancy probabilities in response to increasing anthropogenic pressures (pressures are scaled). C: Means (coloured lines) and CrIs of estimated species richness per site in response to increasing anthropogenic pressure (pressures are scaled). 95% CrIs are shown in light grey and 68% CrIs are shown in dark grey. Panels B and C show predicted occupancy and estimated species richness in response to the combined linear and quadratic effects of cattle, because we found significant responses to both (Extended Data Table 1). We performed a multicollinearity check and found that | r | < 0.7 for all model covariates, allowing us to consider them independently. Silhouettes obtained from PhyloPic (www.phylopic.com).

These findings diverge from earlier studies linking competition with livestock, including cattle, to wildlife declines across the GME and other African savannas^18,21^. However, they align with a long history of coexistence: Maasai pastoralists have grazed cattle alongside high wildlife densities for millennia^22^, and community conservancies build on this legacy. Rather than imposing complete restrictions on mobility as in strict PAs, conservancies implement rotational grazing, opening and closing “grazing blocks” to cattle, attempting to mimic traditional seasonal movements and creating space for wildlife. While some landowners now receive substantial lease payments from tourism, cattle remain central to Maasai livelihoods and identity^23^. Facilitation, where the activities of one species improve forage conditions for another, provides a possible ecological mechanism for this positive cattle-wildlife association^24^. For example, cattle grazing can shorten grasses, promoting fresh regrowth and creating open areas that wild ungulates preferentially use for feeding and vigilance^25^.

Because community-level patterns can mask species-specific vulnerabilities^26^, we also examined whether individual taxa may be more sensitive to existing levels of cattle pressure. Species-specific responses were consistent with that of the community, and all mammal species exhibited positive linear and smaller negative quadratic occupancy associations with cattle presence, with substantial posterior support in most cases (Figure 4, Extended Data Table 3). For some species, such as elephants, we captured higher trap rates in areas without cattle, (i.e., strictly protected Mara Triangle) (Extended Data Figure 3). This greater sensitivity to cattle pressure is supported by occupancy; elephant predicted occupancy declined below baseline at 75% of observed cattle pressure, whereas impala did not decline until 95% (Figure 4, Extended Data Table 2). For some species such as giraffe, observed levels of cattle pressure are not yet detectably decreasing occupancy (Extended Data Table 2). Notably, we captured these patterns during the dry season, when forage competition typically intensifies and facilitation effects are reduced^24,27^, suggesting that positive or neutral responses to cattle are not solely seasonal artefacts.

**Figure 4:**
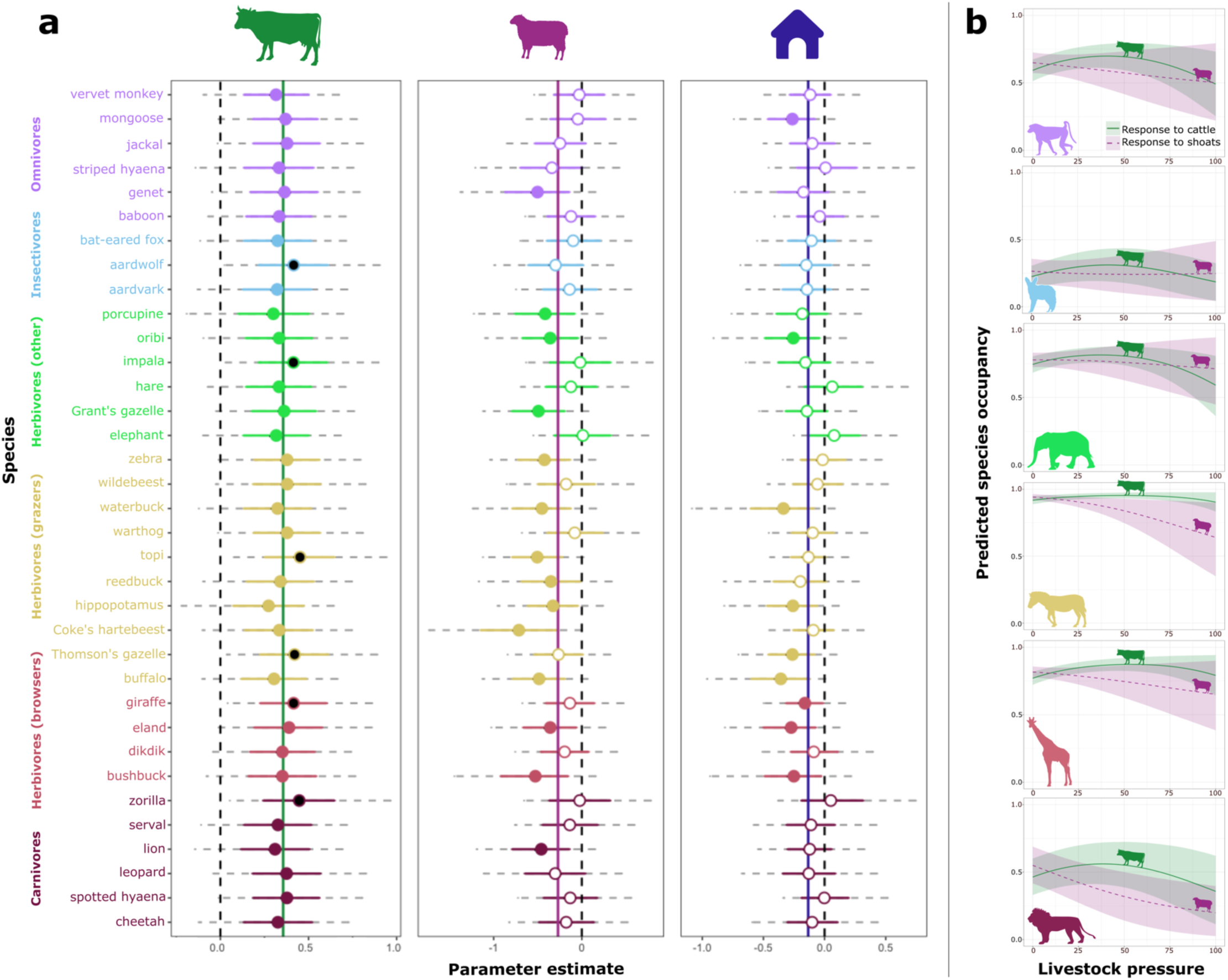
Individual mammal species occupancy responses and predictions along three anthropogenic pressure gradients. A: Model coefficients for the effects of cattle grazing, shoat grazing, and distance to human infrastructure on mammal species occupancy. Solid vertical lines show mean community occupancy estimates. Circles show individual species posterior means with 68% (solid coloured line) and 95% (dashed grey line) Bayesian credible intervals (CrIs). Closed coloured circles indicate estimates with 68% CrI which do not overlap zero, and closed black circles indicate estimates with 95% CrIs which do overlap zero. B: Predicted species occupancy in response to increasing cattle pressure (green) and shoat pressure (magenta). Lines represent means, shown with 68% CrIs. Plots shown for six example species, each belonging to a different functional dietary group: olive baboon (omnivore), bat-eared fox (insectivore), African elephant (herbivore, other (mixed grazer and browser)), plains zebra (herbivore, grazer), giraffe (herbivore, browser), and African lion (carnivore). Silhouettes obtained from PhyloPic www.phylopic.com.

Our study suggests that these conservancies are successfully acting as mixed-use conservation areas, accommodating cattle production, tourism, and biodiversity goals^3^. GME cattle numbers remained relatively stable between 1977 and 2009, though their distribution expanded^18^ and anecdotal reports suggest recent increases. Our results indicate that while current cattle levels generally allow wildlife to persist, further increases in cattle pressure risk disrupting coexistence^28^. Adaptive management will be essential to maintain these wildlife populations sustainably, and quantifying the limits of this coexistence remains a priority. For example, some conservancy actors hope to decrease cattle numbers through herd efficiency: crossbreeding traditional drought-tolerant species (primarily Small East African Zebu) with introduced bulls of higher-yielding species (primarily Sahiwal and Borana). Our findings reinforce emerging evidence in East Africa that wildlife can coexist with cattle at moderate densities^5,29–31^.

## Unmanaged sheep/goat grazing is detrimental

In contrast to cattle, the wildlife community responded negatively to shoat grazing with a posterior mean of -0.27 and high posterior inclusion probability (f = 0.93). We found substantial posterior support for this negative relationship with a 68% credible interval excluding zero and a 95% credible interval marginally overlapping zero (68% CrI = -0.45 to -0.09, 95% CrI = -0.62 to 0.08, Figure 3A, Extended Data Table 1). Community predicted occupancy declined by 25.7% at 50% of observed shoat pressure, and predicted species richness also declined steadily (Figure 3B, 3C, Extended Data Table 4), suggesting that elevated shoat densities may constrain mammal diversity and community occupancy. The more negative impact of shoats than cattle likely reflects their more intensive and destructive foraging strategies^32^, a concern given the rapidly increasing investment in shoats across African savannas^33^. In the GME, shoat numbers more than tripled between 1977 and 2014^23^. This herd diversification is a common adaptation to restricted pastoral mobility, and reflects growing climatic and economic uncertainty. Shoats are more drought-resilient than cattle, reproduce quickly, and sell easily for rapid income^32^. Despite their growing prevalence, shoat grazing remains largely unmanaged within conservancies beyond being broadly restricted to settlement areas.

Although livestock impacts on wildlife have been studied extensively across the GME^5,11,32–34^, species-specific livestock effects are rarely distinguished. ‘Livestock’ is frequently used interchangeably with ‘cattle’ when evaluating management, overlooking the distinct ecological impacts of shoats^18^. While cattle and sheep are preferential grazers and goats are browsers, all three browse in dry periods^35^. Shoats graze more intensively than cattle, and sheep can uproot entire grasses, accelerating desertification^33^. The spread of fast-breeding, disease-resistant Dorper sheep has likely escalated this pressure^32^. Shoats are emerging as a major ecological pressure requiring active management, and we can reasonably expect numbers to rise under worsening climate variability^23^. Uncontrolled growth of shoat populations degrading rangelands was a weakness identified in the group ranch system which preceded conservancy development^36^. Moreover, socio-cultural demands for shoats (e.g., for ceremonies and gifts) have cemented their place in society^37^. In their current unregulated state, shoats present a significant risk to the wildlife that underpins ecotourism, local livelihoods, and the long-term resilience of community-based conservancies.

The negative community-wide effects of shoats extended to individual mammal species (Figure 4, Extended Data Table 2). While all species showed negative occupancy responses, they varied in their strength of decline (Extended Data Table 4). Between 0 and 50% observed shoat pressure, some species’ predicted occupancy declines by over 40% (e.g., hartebeest, buffalo), while others decline by less than 10% (e.g., giraffe, spotted hyaena) (Extended Data Table 4). Although sheep and goats are typically grouped due to mixed herds and identification challenges, this practice may mask differences in ecological roles: goats as key browsers and sheep as more intensive grazers. Wealthier households may increasingly favour sheep over goats^38^, potentially accelerating a shift towards species with greater ecological impact. Disentangling their impacts in future work will be essential for developing effective, adaptive grazing management in wildlife-rich rangelands.

## Infrastructure proximity reduces mammal occupancy

Mammal community occupancy declined with increasing proximity to human infrastructure, including settlements, towns, and bomas (mean = -0.13, f = 0.94). We found substantial posterior support for this negative relationship with a 68% credible interval excluding zero and a 95% credible interval marginally overlapping zero (68% CrI = -0.22, -0.05, 95% CrI = -0.31, 0.04, Figure 3A, Extended Data Table 1). Predicted occupancy declined by 21.4% at 50% of observed pressure, and species richness similarly declined linearly, though with smaller effect sizes than for shoats (Figure 3B, 3C, Extended Data Tables 1, 4). This negative relationship was largely consistent across species, with predicted occupancy declines up to 46% (buffalo) at 50% observed infrastructure pressure (Figure 4A, Extended Data Tables 3, 4). Some species did show marginal positive responses to infrastructure (e.g., hare, elephant), but these effect sizes are small with 68% credible intervals overlapping zero (Figure 4A, Extended Data Table 3). This largely negative pattern likely reflects the ecological consequences of recent sedentarisation and land-use intensification.

Historically, Maasai pastoralists were transhumant people, tracking rainfall and pasture availability in shared landscapes with wildlife^34^. In recent decades, however, land fragmentation driven by privatisation policy has led to widespread subdivision and fencing^32,39^. This shift has curtailed Maasai mobility, making pastoral livelihoods untenable, encouraging permanent settlement, and concentrating infrastructure in formerly open rangelands. Mobility and forage availability for people and livestock were already reduced by their exclusion from strictly protected areas such as MMNR^40^, and this increasing sedentarisation has been linked to wildlife declines through land-use intensification, forage reduction, corridor disruption, and direct displacement^19,23,34,41^. Ongoing land subdivision among the next generation of landowners is likely to accelerate infrastructure spread across the GME. Projections suggest this could cause >40% declines in wildlife abundance and the local loss of key tourism species^34,42^. Sedentarisation also concentrates grazing near homesteads, compounding ecological impacts^32,43,44^. While infrastructure effects on occupancy are currently moderate, continued land-use change points to an increasingly fragmented landscape in which wildlife persistence and conservancy viability may become untenable without proactive spatial planning.

## Discussion

Our findings demonstrate that community conservancies can enable functional coexistence between wildlife and pastoral livelihoods. Maasai landowners retain aspects of traditional pastoralism through rotational cattle grazing, with lease payments providing supplementary income, though recent COVID-19 travel bans exposed instability in overreliance on international tourism^45^. We found broad evidence for cattle-wildlife coexistence, though fine-scale sensitivities suggest that this system cannot sustain indefinite increases in cattle, and the highest hotspots of cattle pressure in our study area may already be negatively impacting some species. More crucially, as pastoralists adapt to a changing climate and shifting economic pressures, the proliferation of shoats risks undercutting the ecological foundations of CBC. Far fewer studies have assessed wildlife interactions with sheep in Kenya compared to research on cattle or livestock generally^32^. However, our results echo findings in northern Kenya that wildlife is most abundant in areas with moderate cattle pressure, but less common near sheep and goats^29^. Many conservancy leases will be renegotiated in coming years and rotational grazing plans are updated regularly, highlighting the need for targeted, livestock-type-specific management to sustain both wildlife populations and resilient livelihoods in mixed-use landscapes.

We found ecological evidence that conservancies are working for wildlife, but the system is imperfect. CBC can perpetuate social inequality, for example due to local elite dominance over land distribution or complex power dynamics^46^. Severe negative social outcomes of CBC have included exploitation and violence^47^. Critically, when CBC governance is locally led rather than externally or top-down, both social and ecological outcomes are fairer and more positive^2,48^. The three conservancies in this study follow a similar governance model where leadership is accountable to a board, whose membership is 50-50 between Maasai landowners and tourism partners^3^. Management plans are developed in consultation with specific groups of landowners (e.g., the grazing committee) then approved by the board^32^. We do not evaluate this governance here, and while this multi-level structure does not guarantee perfect implementation, it does prioritize participatory decision-making which likely enables coexistence with wildlife. Though human-wildlife conflict persists in this area, evidenced by livestock depredation and retaliatory poisonings^49^, lease agreements continue to be extended, suggesting that this CBC model has been beneficial to both local livelihoods and conservation through ecotourism^11^.

Finally, our study highlights the value of integrating AI tools such as our MMC to monitor human-wildlife coexistence. By automating species identification across over two million camera trap images, we generated fine-scale, multispecies data capturing wildlife responses to co-occurring anthropogenic pressures across a complex landscape at unprecedented spatial scales. These tools can dramatically reduce analytical lag and equip local decision-makers with efficient, actionable information^50^. Making these technologies open source, accessible, and usable locally can also strengthen the participatory dimensions of community-based conservation. When implemented ethically and thoughtfully, AI-assisted pipelines can align scientific evidence with locally-led governance^6^. As CBC is increasingly promoted to meet global biodiversity targets, its long-term viability will depend on tools and policies that are responsive to emerging pressures and grounded in timely ecological feedback. Ultimately, this integration supports the conservation of landscapes where humans, livestock, and wildlife can coexist sustainably.

## Methods

### Data collection

We deployed 180 unbaited camera traps (Browning Dark Ops 2017) in an array of 2 km grids spanning four adjacent conservation areas within the Greater Maasai Mara Ecosystem (GME) in Narok County, south-western Kenya (centred at 1°S, 35°E; elevation ∼1700m) (Figure 1) from 5 October 2018 to 29 November 2018. We used 2km grids to ensure substantial geographical coverage of each area, following other well-established camera trapping protocols for large mammal community assessments^51–53^. Annual prescribed burning occurs only in Mara Triangle. The area is predominantly semi-arid open grassland with higher tree and shrub density along drainage lines^33,54,55^. Rain falls across the study area in a gradient, increasing towards the north-west^4,56^, and annual rainfall has a bimodal pattern: a shorter wet season from mid-November to December and a longer one between March and June^57^.

Single cameras were mounted on trees or poles, 50cm above the ground, within a 200m radius from the centre of each grid and serviced monthly. Passive infrared sensors triggered image capture with a one second delay between triggers. To standardise effort, we excluded the first and last four days when not all the cameras were active, yielding a final survey period from 9 October to 25 November 2018. We pooled these data into 12 sampling occasions of four 24-hour periods to improve occupancy model convergence, following an approach commonly adopted with camera trap data^58–60^. We excluded 31 cameras that operated for fewer than three sampling occasions, and two cameras were stolen, resulting in a final dataset comprising 147 camera stations (Figure 1). This effort, 147 stations over 47 days, meets or exceeds thresholds shown to yield robust occupancy estimates for the majority of species in an ecosystem, allowing for moderate to high variation of occupancy and including rare taxa with occupancy estimates <0.25^61^. Conducting the study within the dry season, when wildlife-livestock competition is also likely to be heightened, minimised the influence of seasonal variation in climatic conditions on subsequent analyses^61,62^.

Monitoring human presence was not a goal of this study, but unintentional bycatch of people (primarily livestock herders and tourists) was unavoidable. These camera traps were deployed in 2018, before any of the now-widespread guidelines for camera trapping with regards to human surveillance had been published^63,64^. In line with emerging best-practices^65^, we partnered with a local NGO (WWF-Kenya), secured permissions from all conservancy boards, and regularly communicated study objectives and findings to local stakeholders and rightsholders. No images or metadata relating to human presence were shared beyond the core research team. Where both humans and wildlife appeared in an image, only the animal record was retained. All images were stored securely and handled in compliance with the UK Data Protection Act of 2018.

### Species classification using a deep learning pipeline

#### Model training and architecture

We trained a customised species classifier, the Maasai Mara Classifier (MMC), based on the ConvNeXt-T architecture^66^ (Figure 2). To improve classification accuracy, we used a ConvNeXt-T model pre-trained on the larger ImageNet dataset^67^, a process that has demonstrated significant gains in species classification performance in camera trap images^68^. The training, test, and validation datasets were formed of crops drawn from one image per 5-minute interval across the dataset and manually labelled using the Visual Object Tagging Tool (VoTT) version 2.2.0^69^ to generate a bounding box and add an annotation. Experts labelled crops based on existing familiarity with the ecosystem and a reference manual with clear images of each species, creating a manual library of 82,300 crops from 37,461 images. These manual labels were accuracy checked by randomly sampling 10% of images per species or species group, and any species with poor sampling accuracy in the sample (>3% error rate) were relabelled entirely. The MMC was trained on 53,102 expert manually annotated image crops from 24,646 images, where an individual crop denotes the 2D bounding box encompassing the pixels a single animal occupies in a camera trap image. It was tested on 23,219 crops from 11,120 images, and validated on 5,879 crops from 4,872 images.

To improve classification performance, certain visually similar or rare species were grouped prior to training (e.g., jackal, hare, shoat, hyena_aardwolf) (Supplementary Table 1). Ambiguous images or rare species were excluded, resulting in 49 final classes, including an ‘other’ label for non-target triggers. The MMC was trained for 100 epochs, using a stochastic gradient descent (SGD) (learning rate: 0.005, momentum: 0.9, weight decay: 0.0001) with input crops resized to 224×224 pixels. Training and validation were performed in PyTorch^70^. As is standard for multi-class classification, we used categorical cross-entropy loss to penalise misclassifications. Data were split 9:4:1 into training, testing, validation sets (Figure 2). MMC performance was evaluated using top-1 and per-class accuracy (Supplementary Table 1, Extended Data Figure 2). Top-1 accuracy = 88.5%, precision = 93.8%, recall = 74.4%, and F1 score = 82.

#### Inference pipeline and data processing

We applied a two-step classification pipeline to our full dataset of 2,078,276 camera trap images.

1. *Animal Detection*. First, we used MegaDetector v4^71^, a pre-trained object detection model, to detect and localise animals. We retained all detections with confidence ≥0.65 based on a subset analysis that balanced false positives and recall, yielding 1,930,714 individual animal crops. Detections with bounding boxes <32 pixels in width/height or <4,096 pixels^2^ in area, were excluded due to poor classification performance on very low-resolution crops.
2. *Species Classification*. Second, we applied the MMC to predict a label and confidence score for each crop. For further analysis and to reduce false positives, we applied a minimum confidence threshold to model outputs where both the MegaDetector and classifier were ≥0.9. While this may increase likelihood of false negatives, this risk was mitigated by aggregating data in 4 x 24-hour sampling periods (96 hours), which increased the probability of species detection over time.

#### Post-processing and quality control

Despite applying a 0.90 confidence threshold and accuracy for cattle and shoats being 0.95 and 0.97 respectively, the MMC produced false positive livestock detections in the Mara Triangle, where we know that no grazing is permitted, and that this policy is strictly enforced. As the average number of predicted cows per cattle photo across the three conservancies was 4.5, versus 1.1 in Mara Triangle, we applied a herd-size filter, retaining livestock detections only if at least two individuals were present per image, consistent with regional herding practices. Manual review of 36,203 images from the Mara Triangle in the expert-labelled subset confirmed zero cattle detections, and this filter successfully eliminated false positive livestock detections in Mara Triangle. While illegal night grazing regularly occurs in some regions of the MMNR^14,72^, it has not been often documented in Mara Triangle, which is managed separately. Additionally, we manually re-identified all detections labelled as ‘hyena_aardwolf’ into species-specific labels (spotted hyena, striped hyena, and aardwolf) prior to analysis to reflect their distinct ecological roles.

#### Final dataset and validation

After filtering, the dataset comprised 515,173 images, yielding 1,184,862 MMC-labelled detections across 37 species excluding certain categories including (e.g., ‘other_rodents’), bird groups, and non-target domestic species such as dogs and donkeys. MMC accuracy for most included species was above 80% (Supplementary Table 1). Rodents, with the exception of porcupines, were excluded from the analysis since most camera traps are not optimized for the detection of small mammals, and are less likely to trigger the camera. In addition to validation during training, we compared multi-species occupancy models (MSOMs) outputs derived from expert-labelled images to those from classifier-labelled on the same subset. Outputs were highly congruent (Extended Data Table 5) supporting the use of automated labels for ecological inference.

#### Anthropogenic pressures and environmental covariates

We quantified anthropogenic pressure at each camera trap station using three metrics: proximity to human infrastructure, cattle trap rate, and shoat trap rate (Extended Data Figure 1). Human infrastructure data was based on a polygon data layer created from Google Earth and Bing maps by Klaassen & Broekhuis^73^ and is based on the shortest distance from each camera trap station to human development, including settlements, bomas, towns, dams and agriculture. Cattle and shoat trap rates were calculated separately to reflect distinct ecological impacts, using:

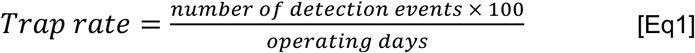

where detection events were identified using classifier labels and defined with a 30-minute independence threshold between individual captures. Camera trap rate has been shown to be a valid index of species density^61,74^. We accounted for a potential nonlinear relationship by including both linear and quadratic terms for livestock rates in the model.

Because the conservancies graze cattle rotationally, our grazing pressure measures may have been biased by the management schedule in some parts of the study area. However, Herrik et al.^75^ collected transect count data across eight months in both the wet and dry seasons in Mara North, and similarly found evidence to support cattle-wildlife coexistence in this area.

Anthropogenic pressure values differed between conservation areas. ANOVA results were significant for all three pressure gradients, indicating that at least one conservation area has a statistically different level of pressure from the rest (Extended Data Figure 1). To compare levels of pressure between each pair of conservation areas, we conducted multiple pairwise t-tests (Welch’s 2-sample t-tests) and applied a Bonferroni correction to p-values to reduce Type I (false positive significance) errors (Extended Data Figure 1).

We also included three environmental covariates hypothesized to affect target species occupancy: proportion of open vegetation, shortest distance to water, and Soil Adjusted Vegetation Index (SAVI), which represents vegetation health and photosynthetic activity. Proportion of open vegetation and distance to water were derived from regional GIS layers^73,76,77^ and calculated within a 500m radius of each camera station. SAVI was computed from Sentinel-2 imagery (10m resolution, 18 November, 2018) using near-infrared (NIR) and visible red (R) bands as:

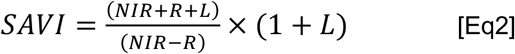

where the soil adjustment factor L = 0.5 was selected for intermediate vegetation density^78^. Mean values were calculated within a 500m radius of each station. SAVI values were generally higher (indicating a larger amount of green vegetation cover) in Mara Triangle, followed by Mara North, Olare-Motorogi, then Naboisho.

To model heterogeneity in detection probability, we obtained data from two variables that are likely to influence the camera trap’s field of view and consequently the detectability of species: tree/shrub density and average grass height directly in front of the camera. These variables were measured in the field during the camera trap survey as follows: tree and shrub density was estimated in the detection field made at each camera station; ten random grass height measurements were made in the detection field at each camera station several times during the survey period and averaged.

### Multispecies occupancy model

We assessed the effects of anthropogenic and environmental covariates on mammal community and species-specific occupancy for 35 species using a Bayesian hierarchical multispecies occupancy model ^79,80^. Detection histories were grouped in 12 sampling occasions of 96 hours (4 x 24 h periods). Prior to modelling, we checked for multicollinearity among covariates; all retained variables had | r | < 0.7. Numerical covariates were standardised (mean = 0, SD = 1). This model was also used to obtain the sum of species at a camera trap site for each iteration of the Bayesian sampling process^60,81^. To estimate species richness and account for undetected species expected in the study area, we used data augmentation with 36 all-zero detection histories, based on the total number of non-rodent terrestrial mammal species expected in the area according to IUCN minus those detected^82^. To examine spatial distribution of wildlife captured across the study area displayed in Extended Data Figure 3, we also calculated trap rates for individual species following [Eq. 1].

We defined occupancy *z_j,I_* as 1 if camera station *j* was occupied by species *i*, and 0 if not.

The occurrence probability (*ψ*) was defined as:

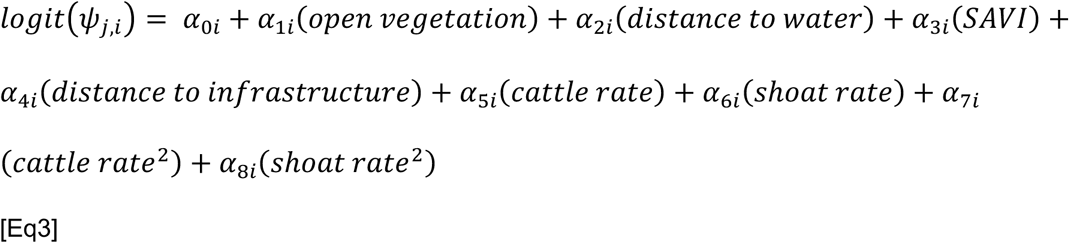

The detection probability (*p*) was defined as:

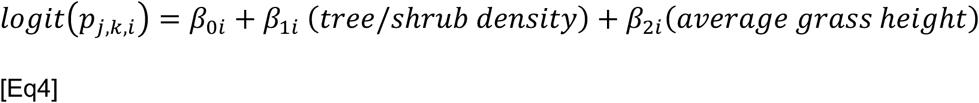

where *log*(*ψ_j,i_*) is the probability that species *i* occurs at the station *j*, and *log(p_j,k,i_*) is the probability of detection for species *i* at station *j* during occasion *k*.

The model was fitted in JAGS v. 4.3.0^83^ via R version 4.0.3^84^ using the *jagsUI* package^85^. We used a burn-in of 50,000 iterations, three chains of 150,000 iterations, and a thinning rate of 10. Convergence was assessed using *R̂* statistics; all parameters had *R̂* ≤ 1.1^86^. We used vague priors for all parameters estimated and conducted a prior sensitivity analysis (Supplementary Table 2). We used the mean of the posterior distribution of each parameter for inference. We interpreted covariate effects as having strong support when 95% credible intervals did not overlap zero, and moderate support when 68% credible intervals (∼ ±1 SD of the mean) did not overlap zero. Re-running the model with separate hyperparameters for herbivores (grazing, browsing, and other herbivores) and non-herbivores (carnivores, omnivores, and insectivores) confirmed that herbivore dominance in detections did not bias community-level responses (Supplementary Tables 1, 3, 4).

To further investigate fine-scale sensitivities to cattle, shoat, and human infrastructure pressure, we obtained predicted occupancy across the full observed range of observed pressures, scaled 0-100. We define the community and each species’ local baseline as predicted occupancy when pressure = 0. Since we found a quadratic occupancy response to cattle pressure, we extracted at what percentage of observed cattle pressure predicted occupancy dips below the local baseline, indicating that cattle pressure is negatively affecting that species’ occupancy (Extended Data Table 2). To quantify linear sensitivities to shoat and human infrastructure pressure, we extracted each species’ predicted occupancy at 50% of observed pressure and calculated the percentage decline in occupancy relative to the baseline (Extended Data Table 4).

## Acknowledgements

This research was funded by WWF-UK as part of the Biome Health Project (www.biomehealthproject.com) and GMM’s BBVA Foundation Frontiers of Knowledge Awards. Camera trap data were collected under E.K.M.’s NACOSTI permit (Ref. no: NACOSTI/P/18/61494/23703) and Kenya Wildlife Service permit (Ref: KWS/BRM/5001). We thank the Mara Triangle, Mara North Conservancy, Naboisho Conservancy, Olare-Motorogi Conservancy, and WWF-Kenya for allowing us to conduct this research and providing rangers for support in the field. Thanks to S. Hatfield for assistance with manual data cleaning. D.J.I. acknowledges funding from UK Research and Innovation Future Leaders Fellowship (Grant ref: MR/W006316/1). HAIP and OP were supported by GMM’s BBVA Frontiers in Knowledge Award. EDM was funded by the European Union (ERC, BIOBANG, 101171602). Views and opinions expressed are however those of the author only and do not necessarily reflect those of the European Union or the European Research Council Executive Agency. Neither the European Union nor the granting authority can be held responsible for them. EDM would also like to thank the KONE Foundation under project #202309134.

## Author CRediT Contributions Statement

Conceptualization: BC, GMM, KEJ

Data curation: EC, HAIP, OP, EKM, MN, LP, AR, FS, LT, AP, GC, SC, TB

Formal analysis: HAIP, EC, OP

Funding acquisition: BC, GMM, KEJ

Investigation (data collection): EKM, DJI

Methodology: GBF, OP, OMA, HAIP, EC, DJI

Project administration: KEJ, GBF, DJI

Resources: KEJ, GMM, PDO

Supervision: KEJ, GBF, DJI, OMA, GJB, MR, RME, EW, EDM

Writing – original draft: EC, HAIP, KEJ

Writing – review & editing: All authors excluding BC and GMM who were deceased before writing began.

Authors after DJI are listed alphabetically up until BC and GMM (deceased) and KEJ (senior author).

## Competing interests

DJI is trustee for The Pangolin Project CIO, working to improve pangolin conservation in Kenya. DJI is a Field Science Co-Chair of the IUCN SSC Pangolin Specialist Group. GJB works part-time for Niantic Spatial UK. KEJ is a trustee of UNEP-WCMC, and a scientific advisor to the Bat Conservation Trust, The Nature Conservancy, and chirrup.ai.

## Notes

### Summary of Updates

Figure 1 updated to reflect most recent submitted version

