## Extended data and supplementary information for "Sustainable cattle management by communities supports African wildlife"

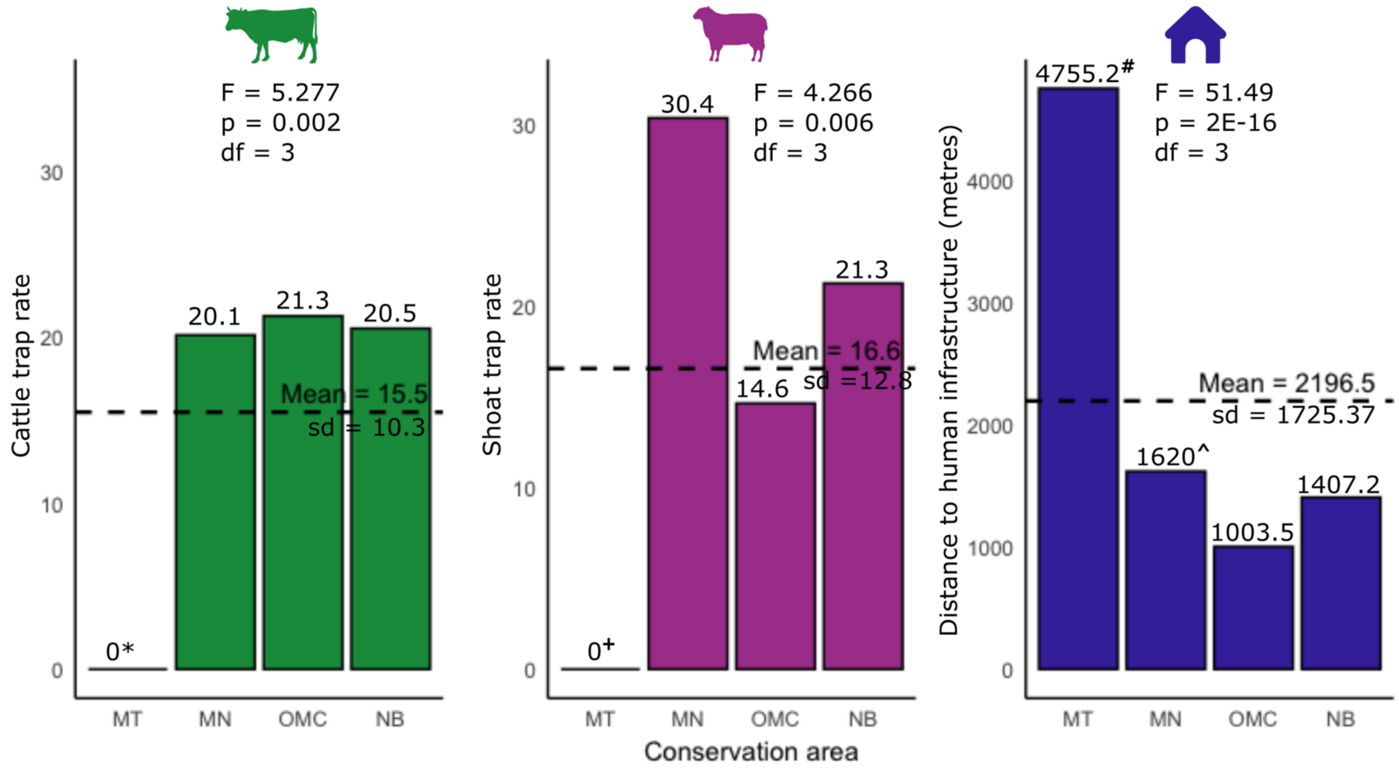


***Extended Data Figure 1: Mean levels of three anthropogenic pressures across camera trap sites in four conservation areas in the Maasai Mara, Kenya.*** *Where MT = Mara Triangle, MN = Mara North Conservancy, OMC = Olare-Motorogi Conservancy, and NB = Naboisho Conservancy. Livestock trap rates were calculated as (number of detection events x 100)/number of camera days) for each camera trap. The distance to human infrastructure pressure is displayed in untransformed units, so higher numbers = lower pressure. ANOVA results were significant for all three pressure gradients; F-statistics and p-values are displayed. Significant results from pairwise t-tests (corrected $p < 0.05$) are denoted with superscript symbology as follows (Methods). *Cattle trap rate in Mara Triangle is significantly lower than in the conservancies (MN t = -3.500, p = 0.008, df = 35; NB t = -4.808, p = 0.0001, df = 44; OMC t = -2.952, p = 0.041, df = 25). ${}^{+}$Shoat trap rate is significantly lower in Mara Triangle than in Mara North (t = -3.480, p = 0.008, df = 35) and Naboisho (t = -3.252, p = 0.013, df = 44). ${}^{\#}$Camera traps in Mara Triangle are significantly further from human infrastructure than camera traps in the conservancies (MN t = 7.341, p = 5.83E-09, df = 55.713; NB t = 8.315, p = 5.87E-10, df = 46.249; OMC t = 9.246, p = 2.19E-1, df = 47.1781), and ^camera traps in Mara North are significantly further from human infrastructure than camera traps in OMC (t = 2.745, p = 0.048, df = 57.957).*

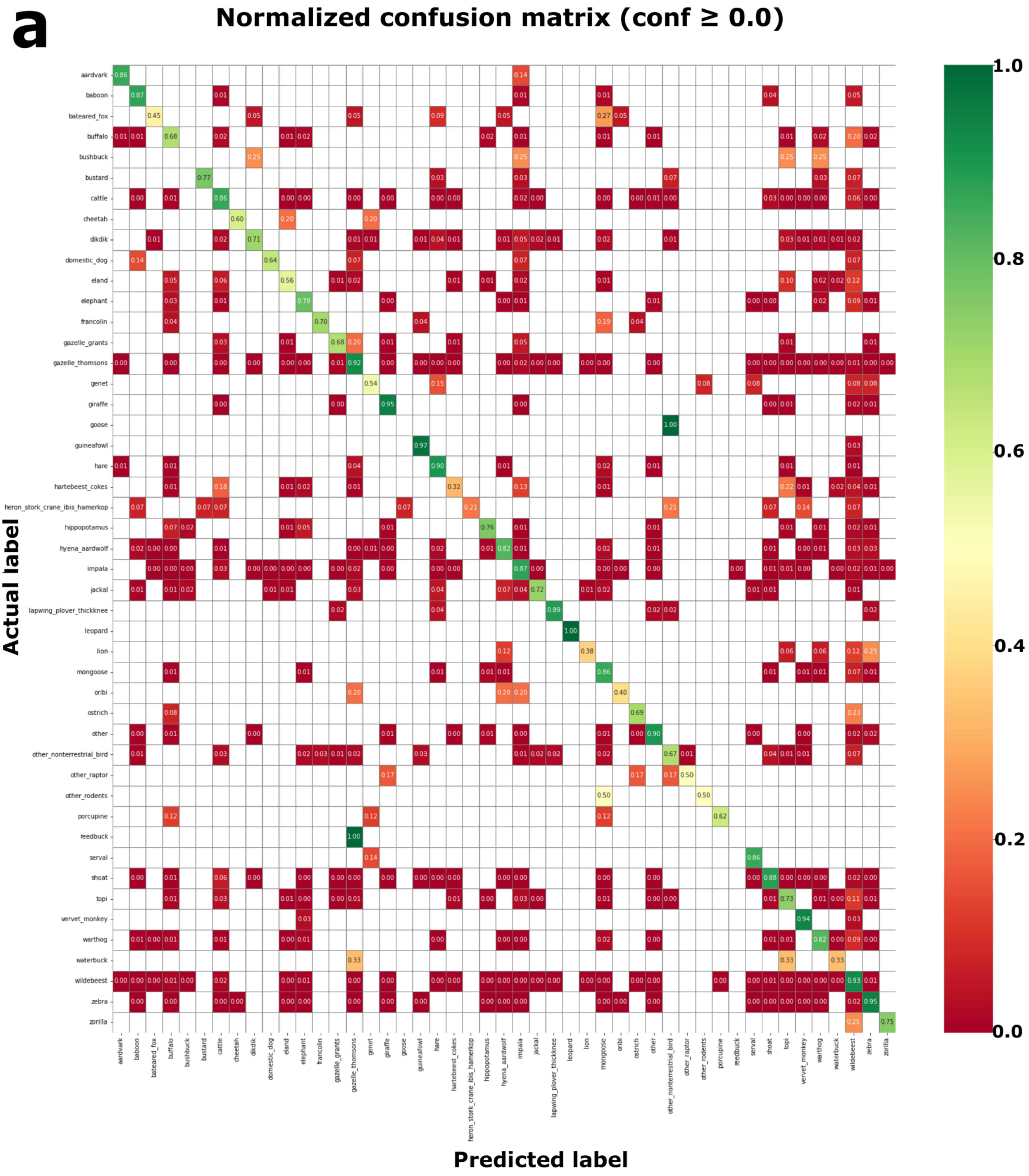
a
Normalized confusion matrix (conf ≥ 0.0)
Actual label
Predicted label
aardvark
baboon
bateared_fox
buffalo
bushbuck
bustard
cattle
cheetah
dikdik
domestic_dog
eland
elephant
francolin
gazelle_grants
gazelle_thomsons
genet
giraffe
goose
guineafowl
hare
hartebeest_cokes
heron_stork_crane_ibis_hamerkop
hippopotamus
hyena_aardwolf
impala
jackal
lapwing_plover_thickknee
leopard
lion
mongoose
oribi
ostrich
other
other_nonterrestrial_bird
other_raptor
other_rodents
porcupine
reedbuck
serval
shoat
topi
vervet_monkey
warthog
waterbuck
wildebeest
zebra
zorilla
1.0
0.8
0.6
0.4
0.2
0.0

**b**

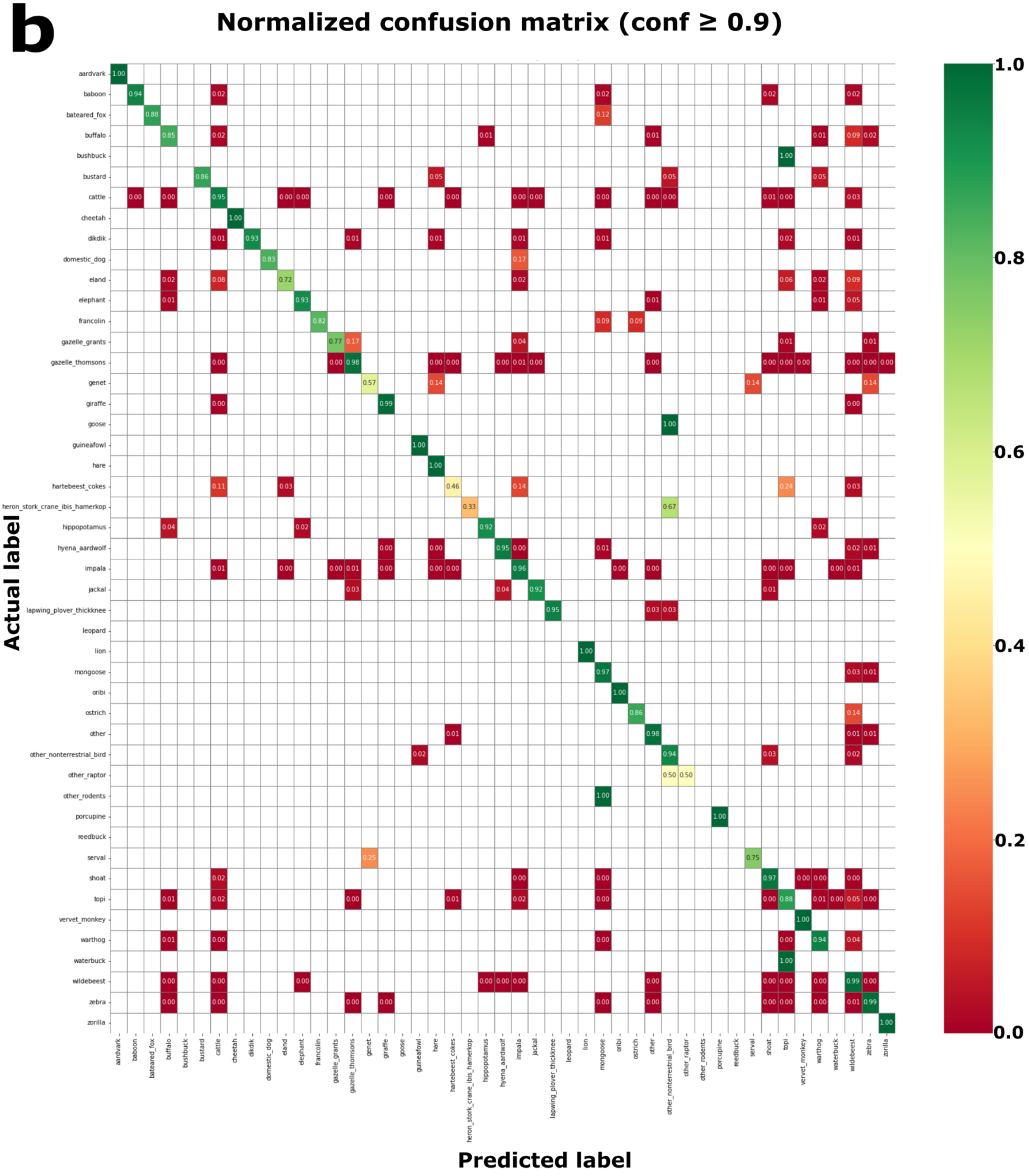
Normalized confusion matrix (conf ≥ 0.9)
Actual label
Predicted label
aardvark
baboon
bateared_fox
buffalo
bushbuck
bustard
cattle
cheetah
dikdik
domestic_dog
eland
elephant
francolin
gazelle_grants
gazelle_thomsons
genet
giraffe
goose
guineafowl
hare
hartebeest_cokes
heron_stork_crane_ibis_hamerkop
hippopotamus
hyena_aardwolf
impala
jackal
lapwing_plover_thickknee
leopard
lion
mongoose
oribi
ostrich
other
other_nonterrestrial_bird
other_raptor
other_rodents
porcupine
reedbuck
serval
shoat
topi
vervet_monkey
warthog
waterbuck
wildebeest
zebra
zorilla
1.0
0.8
0.6
0.4
0.2
0.0

***Extended Data Figure 2: Mara Mammal Classifier performance evaluation: confusion matrix for species classification.*** *Matrices show the proportion of images correctly classified by the MMC across 49 labels including 35 mammal species in our analysis, with A) no confidence threshold applied and B) 90% confidence threshold applied. True species labels are on the y-axis. Cells along the diagonal represent correct classifications, and cells off the diagonal represent common misclassifications. Bird species, rodents and other excluded classes are also displayed here due to possible confusion with mammals. Some species such as bushbuck have no value for B) as low sample size in the test set meant that only false negatives were returned by the classifier.*

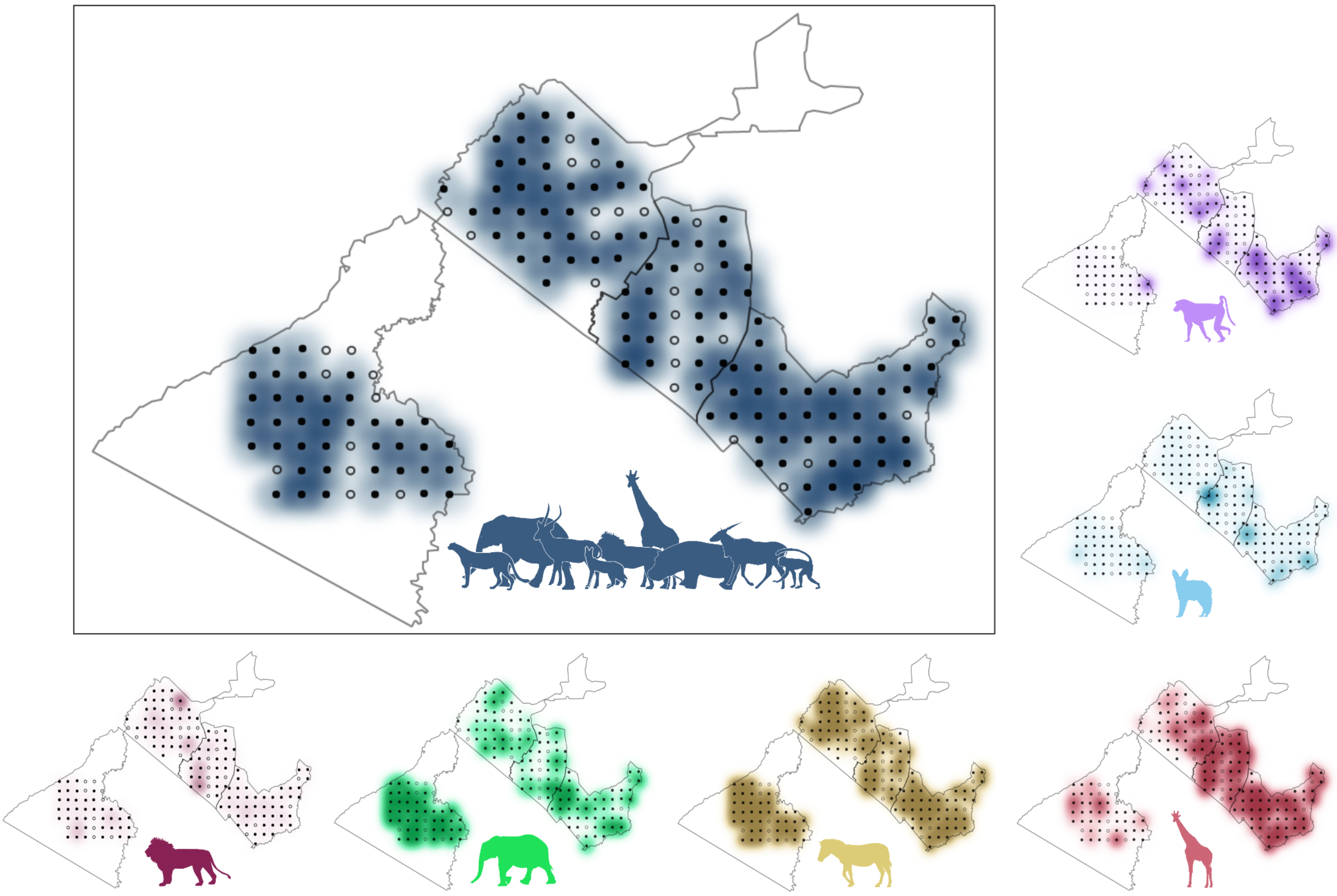

***Extended Data Figure 3: Spatial distribution of wildlife captures in the Maasai Mara, Kenya.*** *Black dots represent locations of 147 camera traps used in our analyses; empty dots represent 33 camera traps excluded due to malfunction or theft. Distributions shown for total species richness (top left box) and trap rate of six example species, each belonging to a different functional dietary group: olive baboon (omnivore), bat-eared fox (insectivore), African elephant (herbivore, other (mixed grazer and browser)), plains zebra (herbivore, grazer), giraffe (herbivore, browser), and African lion (carnivore). Darker colours indicate higher trap rate. Trap rates were calculated as number of detection events x 100/number of full days the camera was operational.* Silhouettes obtained from PhyloPic www.phylopic.com.

***Extended Data Table 1: Community-level effects on occupancy (α) and detection (β) probabilities of mammals in the Maasai Mara, Kenya.*** *Mean, standard deviations, and 95% and 68% Bayesian credible intervals (CrI) for parameter estimates. Parameters with estimates whose 95% and 68% CrI do not overlap zero are highlighted in bold and marked with ** and * respectively.*

| Community-level parameter | Mean | Standard deviation | 95% Bayesian CrI | | 68% Bayesian CrI | | f |
|---|---|---|---|---|---|---|---|
| α1: proportion of open space | -0.065 | 0.114 | -0.295 | 0.156 | -0.176 | 0.045 | 0.721 |
| **α2: shortest distance to water*** | -0.083 | 0.072 | -0.224 | 0.060 | -0.156 | -0.014 | 0.881 |
| α3: SAVI | -0.176 | 0.144 | -0.461 | 0.108 | -0.316 | -0.037 | 0.893 |
| **α4: shortest distance to human infrastructure*** | -0.134 | 0.088 | -0.305 | 0.043 | -0.218 | -0.050 | 0.937 |
| **α5: cattle trap rate**** | 0.356 | 0.147 | 0.062 | 0.645 | 0.211 | 0.500 | 0.991 |
| **A6: shoat trap rate*** | -0.270 | 0.179 | -0.624 | 0.084 | -0.445 | -0.093 | 0.933 |
| **α7: quadratic cattle trap rate**** | -0.090 | 0.043 | -0.174 | -0.004 | -0.132 | -0.048 | 0.979 |
| α8: shoat trap rate$^2$ | 0.027 | 0.061 | -0.096 | 0.150 | -0.032 | 0.086 | 0.680 |
| **B1i: average grass height**** | -0.229 | 0.074 | -0.381 | -0.090 | -0.301 | -0.158 | 0.999 |
| β2i: tree/shrub density | -0.014 | 0.060 | -0.132 | 0.102 | -0.074 | 0.044 | 0.592 |

***Extended Data Table 2: Predicted occupancy for the mammal community and individual species with respect to cattle pressure in the Maasai Mara, Kenya.*** *To further consider species' sensitivity to cattle pressure and quantify this quadratic relationship, we report the local baseline predicted occupancy (predicted occupancy when cattle pressure= 0). For most species, predicted occupancy increases with cattle pressure up to a certain point, after which it begins to decline. Here we also report at what percentage of observed cattle pressure predicted occupancy dips below the local baseline, indicating that cattle pressure is negatively affecting that species' occupancy. For some species, predicted occupancy does not decrease past their local baseline at levels of cattle pressure observed in this study, so we report NA.*

| Species | Local_baseline | %Cattle_decrease |
|---|---|---|
| community | 0.530 | 92 |
| aardvark | 0.349 | 82 |
| aardwolf | 0.197 | NA |
| baboon | 0.593 | 83 |
| bateared_fox | 0.225 | 87 |
| buffalo | 0.571 | 84 |
| bushbuck | 0.597 | 93 |
| cheetah | 0.146 | 94 |
| dikdik | 0.394 | 92 |
| eland | 0.748 | 86 |
| elephant | 0.745 | 75 |
| gazelle_grants | 0.564 | 88 |
| gazelle_thomsons | 0.877 | NA |
| genet | 0.223 | NA |
| giraffe | 0.771 | NA |
| hare | 0.685 | 79 |
| hartebeest | 0.614 | 86 |
| hippopotamus | 0.247 | 73 |

| | | |
|---|---|---|
| hyena_spotted | 0.834 | 89 |
| hyena_striped | 0.413 | 89 |
| impala | 0.917 | 95 |
| jackal | 0.694 | 89 |
| leopard | 0.227 | NA |
| lion | 0.463 | 79 |
| mongoose | 0.565 | 93 |
| oribi | 0.168 | 95 |
| porcupine | 0.337 | 82 |
| reedbuck | 0.090 | NA |
| serval | 0.692 | 80 |
| topi | 0.709 | NA |
| vervet_monkey | 0.303 | 87 |
| warthog | 0.897 | 89 |
| waterbuck | 0.315 | 85 |
| wildebeest | 0.974 | 88 |
| zebra | 0.917 | 95 |
| zorilla | 0.283 | NA |

***Extended Data Table 3: Species-level responses of mammals to anthropogenic pressures in the Maasai Mara, Kenya.*** *Mean, standard deviation, and 95% and 68% Bayesian credible intervals (CrI) for the estimates of individual species responses to three anthropogenic pressure variables: human infrastructure, cattle presence, and shoat presence. Estimates with 68% CrIs that do not include zero are in* ***bold*** *and denoted with *.*

| | **Human infrastructure** | | | | | | **Cattle** | | | | | | **Quadratic cattle** | | | | | | **Shoats** | | | | | |
|---|---|---|---|---|---|---|---|---|---|---|---|---|---|---|---|---|---|---|---|---|---|---|---|---|
| **Species label** | **Mean** | **SD** | **95% CrI** | | **68% CrI** | | **Mean** | **SD** | **95% CrI** | | **68% CrI** | | **Mean** | **SD** | **95% CrI** | | **68% CrI** | | **Mean** | **SD** | **95% CrI** | | **68% CrI** | |
| aardvark | -0.143 | 0.238 | -0.640 | 0.357 | -0.345 | 0.052 | **0.322*** | 0.214 | -0.129 | 0.721 | **0.127** | **0.516** | **-0.103*** | 0.060 | -0.229 | 0.009 | **-0.155** | **-0.049** | -0.139 | 0.328 | -0.746 | 0.561 | -0.442 | 0.171 |
| aardwolf | -0.149 | 0.252 | -0.688 | 0.366 | -0.357 | 0.057 | **0.416**** | 0.220 | **0.024** | **0.905** | **0.216** | **0.610** | **-0.088*** | 0.060 | -0.199 | 0.042 | **-0.142** | **-0.037** | -0.298 | 0.336 | -0.993 | 0.358 | -0.608 | 0.014 |
| baboon | -0.040 | 0.210 | -0.410 | 0.439 | -0.225 | 0.158 | **0.334*** | 0.196 | -0.069 | 0.712 | **0.146** | **0.518** | **-0.094*** | 0.055 | -0.201 | 0.020 | **-0.146** | **-0.044** | -0.122 | 0.282 | -0.640 | 0.473 | -0.392 | 0.153 |
| bateared_fox | -0.107 | 0.227 | -0.551 | 0.382 | -0.299 | 0.087 | **0.325*** | 0.207 | -0.114 | 0.714 | **0.133** | **0.516** | **-0.100*** | 0.058 | -0.219 | 0.011 | **-0.152** | **-0.048** | -0.098 | 0.319 | -0.697 | 0.563 | -0.398 | 0.208 |
| buffalo | **-0.358*** | 0.251 | -0.961 | 0.007 | **-0.599** | **-0.126** | **0.304*** | 0.194 | -0.103 | 0.665 | **0.117** | **0.488** | **-0.086*** | 0.052 | -0.184 | 0.024 | **-0.135** | **-0.039** | **-0.485*** | 0.303 | -1.127 | 0.067 | **-0.786** | **-0.190** |
| bushbuck | **-0.251*** | 0.278 | -0.936 | 0.218 | **-0.491** | **-0.028** | **0.352*** | 0.212 | -0.079 | 0.766 | **0.159** | **0.544** | **-0.084*** | 0.058 | -0.191 | 0.042 | **-0.137** | **-0.035** | **-0.527*** | 0.407 | -1.444 | 0.158 | **-0.920** | **-0.153** |
| cheetah | -0.102 | 0.246 | -0.591 | 0.436 | -0.305 | 0.107 | **0.326*** | 0.215 | -0.123 | 0.728 | **0.133** | **0.521** | **-0.097*** | 0.059 | -0.217 | 0.017 | **-0.150** | **-0.045** | -0.177 | 0.337 | -0.825 | 0.521 | -0.486 | 0.139 |
| dikdik | -0.088 | 0.219 | -0.508 | 0.396 | -0.276 | 0.109 | **0.352*** | 0.196 | -0.042 | 0.743 | **0.167** | **0.536** | **-0.093*** | 0.055 | -0.201 | 0.020 | **-0.145** | **-0.043** | -0.195 | 0.289 | -0.755 | 0.401 | -0.467 | 0.078 |
| eland | **-0.273*** | 0.237 | -0.814 | 0.123 | **-0.501** | **-0.070** | **0.389*** | 0.217 | -0.010 | 0.859 | **0.192** | **0.578** | **-0.097*** | 0.061 | -0.219 | 0.025 | **-0.152** | **-0.045** | **-0.358*** | 0.324 | -1.026 | 0.268 | **-0.666** | **-0.060** |
| elephant | 0.078 | 0.219 | -0.251 | 0.589 | -0.128 | 0.289 | **0.317*** | 0.197 | -0.098 | 0.684 | **0.130** | **0.502** | **-0.100*** | 0.054 | -0.210 | 0.006 | **-0.151** | **-0.050** | 0.012 | 0.335 | -0.549 | 0.756 | -0.312 | 0.329 |
| gazelle_grants | -0.144 | 0.192 | -0.534 | 0.261 | -0.318 | 0.023 | **0.362*** | 0.195 | -0.024 | 0.759 | **0.178** | **0.544** | **-0.095*** | 0.054 | -0.201 | 0.015 | **-0.146** | **-0.046** | **-0.492*** | 0.303 | -1.119 | 0.075 | **-0.792** | **-0.192** |
| gazelle_thomsons | **-0.263*** | 0.202 | -0.704 | 0.098 | **-0.459** | **-0.082** | **0.421**** | 0.212 | **0.036** | **0.891** | **0.226** | **0.612** | **-0.074*** | 0.060 | -0.176 | 0.064 | **-0.129** | **-0.023** | -0.265 | 0.291 | -0.841 | 0.324 | -0.539 | 0.013 |
| genet | -0.172 | 0.253 | -0.734 | 0.329 | -0.386 | 0.032 | **0.364*** | 0.211 | -0.048 | 0.791 | **0.170** | **0.553** | **-0.090*** | 0.059 | -0.202 | 0.032 | **-0.144** | **-0.039** | **-0.502*** | 0.390 | -1.377 | 0.155 | **-0.872** | **-0.140** |
| giraffe | **-0.161*** | 0.165 | -0.497 | 0.170 | **-0.315** | **-0.013** | **0.415**** | 0.204 | **0.041** | **0.861** | **0.225** | **0.603** | **-0.076*** | 0.059 | -0.178 | 0.060 | **-0.130** | **-0.024** | -0.135 | 0.290 | -0.674 | 0.475 | -0.410 | 0.143 |
| hare | 0.063 | 0.257 | -0.307 | 0.682 | -0.164 | 0.306 | **0.333*** | 0.197 | -0.072 | 0.711 | **0.145** | **0.516** | **-0.099*** | 0.054 | -0.206 | 0.008 | **-0.149** | **-0.049** | -0.123 | 0.307 | -0.691 | 0.529 | -0.411 | 0.169 |
| hartebeest_cokes | -0.092 | 0.186 | -0.452 | 0.320 | -0.256 | 0.075 | **0.334*** | 0.212 | -0.101 | 0.747 | **0.138** | **0.525** | **-0.089*** | 0.056 | -0.195 | 0.030 | **-0.142** | **-0.040** | **-0.714*** | 0.450 | -1.733 | 0.001 | **-1.149** | **-0.281** |
| hippopotamus | **-0.259*** | 0.234 | -0.820 | 0.117 | **-0.471** | **-0.060** | **0.273*** | 0.216 | -0.220 | 0.644 | **0.072** | **0.474** | **-0.105*** | 0.057 | -0.227 | 0.000 | **-0.156** | **-0.052** | **-0.327*** | 0.300 | -0.955 | 0.241 | **-0.614** | **-0.043** |
| hyena_spotted | -0.002 | 0.215 | -0.337 | 0.517 | -0.191 | 0.197 | **0.378*** | 0.205 | -0.010 | 0.806 | **0.185** | **0.562** | **-0.088*** | 0.057 | -0.192 | 0.035 | **-0.140** | **-0.038** | -0.132 | 0.322 | -0.723 | 0.562 | -0.433 | 0.175 |
| hyena_striped | 0.008 | 0.293 | -0.457 | 0.732 | -0.227 | 0.264 | **0.333*** | 0.223 | -0.141 | 0.756 | **0.134** | **0.529** | **-0.093*** | 0.061 | -0.211 | 0.029 | **-0.147** | **-0.041** | -0.339 | 0.401 | -1.207 | 0.427 | -0.699 | 0.011 |
| impala | -0.156 | 0.249 | -0.644 | 0.396 | -0.379 | 0.048 | **0.414**** | 0.217 | **0.029** | **0.898** | **0.216** | **0.606** | **-0.081*** | 0.060 | -0.188 | 0.053 | **-0.136** | **-0.030** | -0.019 | 0.367 | -0.631 | 0.806 | -0.366 | 0.327 |

| | | | | | | | | | | | | | | | | | | | | | | | | |
|---|---|---|---|---|---|---|---|---|---|---|---|---|---|---|---|---|---|---|---|---|---|---|---|---|
| jackal | -0.100 | 0.211 | -0.499 | 0.373 | -0.283 | 0.084 | **0.378*** | 0.208 | -0.016 | 0.810 | **0.186** | **0.565** | **-0.092*** | 0.056 | -0.199 | 0.024 | **-0.145** | **-0.042** | -0.247 | 0.309 | -0.849 | 0.388 | -0.534 | 0.044 |
| leopard | -0.129 | 0.265 | -0.676 | 0.430 | -0.342 | 0.089 | **0.376*** | 0.215 | -0.040 | 0.827 | **0.182** | **0.568** | **-0.088*** | 0.060 | -0.201 | 0.040 | **-0.142** | **-0.037** | -0.302 | 0.384 | -1.112 | 0.455 | -0.647 | 0.040 |
| lion | -0.122 | 0.209 | -0.544 | 0.326 | -0.302 | 0.058 | **0.310*** | 0.211 | -0.144 | 0.694 | **0.116** | **0.505** | **-0.098*** | 0.059 | -0.218 | 0.017 | **-0.152** | **-0.046** | **-0.460*** | 0.340 | -1.192 | 0.153 | **-0.792** | **-0.138** |
| mongoose | **-0.263*** | 0.207 | -0.741 | 0.082 | **-0.464** | **-0.079** | **0.370*** | 0.197 | -0.011 | 0.775 | **0.185** | **0.552** | **-0.089*** | 0.056 | -0.194 | 0.031 | **-0.141** | **-0.041** | -0.045 | 0.317 | -0.627 | 0.611 | -0.359 | 0.269 |
| oribi | **-0.256*** | 0.266 | -0.908 | 0.176 | **-0.488** | **-0.040** | **0.333*** | 0.204 | -0.096 | 0.721 | **0.143** | **0.522** | **-0.095*** | 0.057 | -0.211 | 0.017 | **-0.147** | **-0.044** | **-0.359*** | 0.340 | -1.097 | 0.275 | **-0.676** | **-0.042** |
| porcupine | -0.182 | 0.255 | -0.760 | 0.299 | -0.394 | 0.023 | **0.300*** | 0.223 | -0.191 | 0.698 | **0.102** | **0.502** | **-0.097*** | 0.060 | -0.218 | 0.019 | **-0.150** | **-0.045** | **-0.416*** | 0.365 | -1.224 | 0.239 | **-0.759** | **-0.080** |
| reedbuck | -0.199 | 0.265 | -0.823 | 0.281 | -0.418 | 0.013 | **0.340*** | 0.212 | -0.098 | 0.747 | **0.149** | **0.531** | **-0.095*** | 0.059 | -0.213 | 0.020 | **-0.148** | **-0.044** | **-0.351*** | 0.371 | -1.168 | 0.341 | **-0.684** | **-0.015** |
| serval | -0.111 | 0.240 | -0.580 | 0.425 | -0.310 | 0.085 | **0.327*** | 0.207 | -0.109 | 0.719 | **0.134** | **0.517** | **-0.093*** | 0.058 | -0.205 | 0.028 | **-0.146** | **-0.042** | -0.136 | 0.341 | -0.758 | 0.599 | -0.448 | 0.182 |
| topi | -0.132 | 0.156 | -0.446 | 0.195 | -0.275 | 0.011 | **0.452**** | 0.215 | **0.076** | **0.943** | **0.252** | **0.648** | **-0.071*** | 0.064 | -0.176 | 0.084 | **-0.127** | **-0.018** | **-0.506*** | 0.291 | -1.120 | 0.020 | **-0.791** | **-0.221** |
| vervet_monkey | -0.118 | 0.189 | -0.493 | 0.285 | -0.285 | 0.051 | **0.316*** | 0.193 | -0.096 | 0.674 | **0.134** | **0.500** | **-0.092*** | 0.053 | -0.194 | 0.015 | **-0.142** | **-0.044** | -0.029 | 0.292 | -0.540 | 0.592 | -0.314 | 0.256 |
| warthog | -0.097 | 0.197 | -0.455 | 0.350 | -0.270 | 0.076 | **0.379*** | 0.205 | -0.013 | 0.809 | **0.188** | **0.566** | **-0.084*** | 0.060 | -0.192 | 0.048 | **-0.139** | **-0.034** | -0.081 | 0.333 | -0.679 | 0.646 | -0.390 | 0.238 |
| waterbuck | **-0.337*** | 0.296 | -1.084 | 0.084 | **-0.605** | **-0.088** | **0.324*** | 0.208 | -0.122 | 0.711 | **0.133** | **0.514** | **-0.100*** | 0.058 | -0.219 | 0.012 | **-0.152** | **-0.048** | **-0.452*** | 0.352 | -1.237 | 0.163 | **-0.789** | **-0.125** |
| wildebeest | -0.058 | 0.240 | -0.466 | 0.518 | -0.259 | 0.145 | **0.380*** | 0.212 | -0.026 | 0.824 | **0.185** | **0.568** | **-0.083*** | 0.060 | -0.191 | 0.049 | **-0.137** | **-0.032** | -0.178 | 0.356 | -0.850 | 0.588 | -0.501 | 0.150 |
| zebra | -0.016 | 0.204 | -0.341 | 0.468 | -0.196 | 0.173 | **0.378*** | 0.201 | -0.010 | 0.796 | **0.192** | **0.562** | **-0.077*** | 0.059 | -0.180 | 0.055 | **-0.131** | **-0.026** | **-0.424*** | 0.304 | -1.041 | 0.155 | **-0.724** | **-0.131** |
| zorilla | 0.051 | 0.286 | -0.380 | 0.746 | -0.190 | 0.317 | **0.447** | 0.226 | **0.054** | **0.966** | **0.243** | **0.648** | **-0.068** | 0.065 | -0.176 | 0.089 | **-0.126** | **-0.014** | -0.024 | 0.363 | -0.654 | 0.784 | -0.372 | 0.323 |

***Extended Data Table 4: Predicted occupancy for the mammal community and individual species with respect to shoat and human infrastructure pressure in the Maasai Mara, Kenya.*** *To quantify the different sensitivities of mammal species to shoat and human infrastructure pressure, we report the local baseline predicted occupancy (predicted occupancy when pressure = 0). We then report each species' predicted occupancy at 50% of observed pressure, and calculate the difference between the two. Finally we report that difference as a percentage decline in occupancy from the local baseline.*

| | Shoats | | | | Human infrastructure | | | |
|---|---|---|---|---|---|---|---|---|
| **Species** | **Local_baseline** | **50%_pressure** | **Difference** | **%_decline** | **Local_baseline** | **50%_pressure** | **Difference** | **%_decline** |
| community | 0.606 | 0.450 | 0.156 | 25.691 | 0.602 | 0.473 | 0.129 | 21.396 |
| aardvark | 0.400 | 0.343 | 0.057 | 14.241 | 0.410 | 0.311 | 0.098 | 23.963 |
| aardwolf | 0.253 | 0.167 | 0.086 | 34.059 | 0.251 | 0.184 | 0.067 | 26.690 |
| baboon | 0.649 | 0.575 | 0.075 | 11.487 | 0.644 | 0.595 | 0.049 | 7.552 |
| bateared_fox | 0.265 | 0.241 | 0.024 | 9.028 | 0.273 | 0.219 | 0.053 | 19.534 |
| buffalo | 0.659 | 0.390 | 0.269 | 40.783 | 0.670 | 0.356 | 0.314 | 46.853 |
| bushbuck | 0.682 | 0.427 | 0.255 | 37.389 | 0.675 | 0.475 | 0.200 | 29.615 |
| cheetah | 0.177 | 0.143 | 0.034 | 19.105 | 0.178 | 0.150 | 0.028 | 15.529 |
| dikdik | 0.462 | 0.361 | 0.101 | 21.842 | 0.458 | 0.385 | 0.074 | 16.053 |
| eland | 0.809 | 0.643 | 0.166 | 20.534 | 0.816 | 0.603 | 0.213 | 26.100 |
| elephant | 0.778 | 0.761 | 0.018 | 2.269 | 0.768 | 0.799 | -0.031 | -4.090 |
| gazelle_grants | 0.661 | 0.388 | 0.273 | 41.301 | 0.638 | 0.500 | 0.138 | 21.629 |
| gazelle_thomsons | 0.909 | 0.827 | 0.082 | 9.022 | 0.915 | 0.777 | 0.138 | 15.070 |
| genet | 0.296 | 0.144 | 0.152 | 51.370 | 0.281 | 0.196 | 0.085 | 30.197 |
| giraffe | 0.817 | 0.747 | 0.070 | 8.547 | 0.825 | 0.705 | 0.120 | 14.598 |
| hare | 0.735 | 0.660 | 0.075 | 10.153 | 0.715 | 0.735 | -0.020 | -2.793 |
| hartebeest | 0.716 | 0.366 | 0.350 | 48.868 | 0.669 | 0.585 | 0.084 | 12.544 |
| hippopotamus | 0.307 | 0.187 | 0.120 | 39.093 | 0.318 | 0.166 | 0.152 | 47.777 |
| hyena_spotted | 0.868 | 0.808 | 0.06 | 6.948 | 0.861 | 0.843 | 0.018 | 2.132 |

| | | | | | | | | |
|---|---|---|---|---|---|---|---|---|
| hyena_striped | 0.471 | 0.347 | 0.124 | 26.385 | 0.447 | 0.453 | -0.006 | -1.375 |
| impala | 0.933 | 0.915 | 0.019 | 1.994 | 0.939 | 0.870 | 0.069 | 7.387 |
| jackal | 0.756 | 0.623 | 0.133 | 17.647 | 0.750 | 0.655 | 0.095 | 12.709 |
| leopard | 0.270 | 0.197 | 0.074 | 27.276 | 0.267 | 0.217 | 0.050 | 18.787 |
| lion | 0.550 | 0.321 | 0.229 | 41.643 | 0.526 | 0.420 | 0.106 | 20.131 |
| mongoose | 0.620 | 0.585 | 0.035 | 5.593 | 0.658 | 0.417 | 0.241 | 36.649 |
| oribi | 0.218 | 0.126 | 0.091 | 41.883 | 0.224 | 0.121 | 0.103 | 45.878 |
| porcupine | 0.408 | 0.242 | 0.166 | 40.683 | 0.400 | 0.281 | 0.119 | 29.690 |
| reedbuck | 0.117 | 0.071 | 0.046 | 39.265 | 0.117 | 0.077 | 0.04 | 34.035 |
| serval | 0.738 | 0.663 | 0.075 | 10.174 | 0.742 | 0.643 | 0.099 | 13.284 |
| topi | 0.793 | 0.533 | 0.26 | 32.809 | 0.774 | 0.662 | 0.111 | 14.386 |
| vervet_monkey | 0.346 | 0.342 | 0.004 | 1.160 | 0.363 | 0.278 | 0.086 | 23.561 |
| warthog | 0.918 | 0.885 | 0.033 | 3.564 | 0.920 | 0.873 | 0.047 | 5.091 |
| waterbuck | 0.398 | 0.211 | 0.187 | 46.986 | 0.410 | 0.196 | 0.215 | 52.264 |
| wildebeest | 0.980 | 0.962 | 0.018 | 1.837 | 0.979 | 0.968 | 0.011 | 1.108 |
| zebra | 0.942 | 0.804 | 0.102 | 10.793 | 0.932 | 0.918 | 0.014 | 1.483 |
| zorilla | 0.325 | 0.328 | -0.004 | -1.132 | 0.319 | 0.364 | -0.045 | -14.122 |

***Extended Data Table 5: Expert-labelled data subset exploration.*** *To further confirm the accuracy of the AI-labelled data output from the Maasai Mara Classifier (MMC), and ensure that this accuracy carried over to downstream occupancy results, we ran the same occupancy model on the manually tagged subset of images. All of the model estimates and credible intervals (CrI) from the expert-labelled subset show a similar pattern to our main results, supporting the use of the MMC.*

| Community-level hyperparameter | Mean | Standard deviation | 95% Bayesian CrI | | 68% Bayesian CrI | |
|---|---|---|---|---|---|---|
| α1: proportion of open space | -0.17 | 0.12 | -0.41 | 0.07 | -0.29 | -0.05 |
| α2: shortest distance to water | -0.07 | 0.07 | -0.22 | 0.07 | -0.15 | 0.00 |
| α3: SAVI | -0.18 | 0.15 | -0.47 | 0.12 | -0.33 | -0.03 |
| α4: shortest distance to human infrastructure | -0.12 | 0.08 | -0.28 | 0.04 | -0.20 | -0.04 |
| α5: cattle trap rate | 0.23 | 0.14 | -0.05 | 0.50 | 0.09 | 0.37 |
| α6: shoat trap rate | -0.35 | 0.17 | -0.70 | -0.02 | -0.53 | -0.18 |
| α7: cattle trap rate$^2$ | -0.08 | 0.04 | -0.16 | 0.01 | -0.12 | -0.04 |
| α8: shoat trap rate$^2$ | 0.01 | 0.06 | -0.10 | 0.12 | -0.05 | 0.07 |
| β1i: average grass height | -0.25 | 0.09 | -0.43 | -0.08 | -0.34 | -0.16 |
| β2i: tree/shrub density | -0.01 | 0.08 | -0.17 | 0.15 | -0.09 | 0.07 |

### Supplementary Information

***Supplementary Table 1: Mammal species included in this study, with associated AI classifier label, trophic classification and classifier accuracy.*** *A total of 37 labels (35 wildlife, 2 livestock) from the Maasai Mara Classifier (MMC) were used for analysis; some labels include multiple species due to morphological similarity which made identification during the expert labelling process difficult (e.g. jackal) or low classifier resolution (hare, mongoose). Although striped hyaena, spotted hyaena, and aardwolf were grouped during classifier training, these were post-hoc separated by expert labelling due to ecological differences. In total, 42 mammal species were represented. Trophic classifications were based on MammalDIET[87], assigning species to Carnivore, Herbivore, or Omnivore. Carnivores scoring '1' for Insectivore and '0' for MammalEater were reclassified as Insectivore. Herbivores scoring '1' for eating woody vegetation were reclassified as Browsers, those scoring '1' for consumption of herbaceous plants were reclassified as Grazers, and those scoring '1' for both as Other Herbivore. Porcupine, lacking clear scores for either, was also classified as Other Herbivore. For multi-species labels, the broadest trophic category was assigned. Adjustments were made based on additional literature: buffalo as Grazer[88]; Grant's gazelle as Other Herbivore[89] and bat-eared fox as Insectivore, reflecting dietary specialization and morphological traits[90]. MMC accuracy is reported with a 90% confidence threshold applied. Leopard and reedbuck had only one record each in the test set without thresholding, and no records at the 90% threshold. Therefore, their accuracy cannot be reported here. Bushbuck had only one record in the test set at the 90% threshold, but the classifier returned a false negative hence the accuracy of 0.00.*

| MMC label | Common name | Species | Trophic classification | MMC accuracy |
|---|---|---|---|---|
| aardvark | Aardvark | *Orycteropus afer* | Insectivore | 1.00 |
| hyena_aardwolf | Aardwolf | *Proteles cristata* | Insectivore | 0.95 |
| baboon | Baboon | *Papio anubis* | Omnivore | 0.94 |
| bateared_fox | Bat-eared fox | *Otocyon megalotis* | Insectivore | 0.88 |
| buffalo | Buffalo | *Syncerus caffer* | Grazer | 0.85 |
| bushbuck | Bushbuck | *Tragelaphus scriptus* | Browser | 0.00 |
| cheetah | Cheetah | *Acinonyx jubatus* | Carnivore | 1.00 |
| dikdik | Dikdik | *Madoqua kirkii* | Browser | 0.93 |
| eland | Eland | *Tragelaphus oryx* | Browser | 0.72 |
| elephant | Elephant | *Loxodonta africana* | Other_herbivore | 0.93 |
| genet | Genet | *Genetta genetta* | Omnivore | 0.57 |
| giraffe | Giraffe | *Giraffa camelopardalis* | Browser | 0.99 |
| gazelle_grants | Grant's gazelle | *Nanger granti* | Other_herbivore | 0.77 |
| hare | Hare | *Lepus capensis* | Other_herbivore * | 1.00 |
| hare | Springhare | *Pedetes capensis* | Other_herbivore * | 1.00 |
| hartebeest_cokes | Hartebeest | *Alcelaphus buselaphus* | Grazer | 0.46 |
| hippopotamus | Hippopotamus | *Hippopotamus amphibius* | Grazer | 0.92 |
| impala | Impala | *Aepyceros melampus* | Other_herbivore | 0.96 |

| | | | | |
|---|---|---|---|---|
| jackal | Black-backed jackal | *Canis mesomelas* | Omnivore | 0.92 |
| jackal | Side-striped jackal | *Canis adustus* | Omnivore | 0.92 |
| leopard | Leopard | *Panthera pardus* | Carnivore | NA |
| lion | Lion | *Panthera leo* | Carnivore | 1.00 |
| mongoose | Banded mongoose | *Mungos mungo* | Omnivore ** | 0.97 |
| mongoose | Dwarf mongoose | *Helogale parvula* | Omnivore ** | 0.97 |
| mongoose | Egyptian mongoose | *Herpestes ichneumon* | Omnivore ** | 0.97 |
| mongoose | Marsh mongoose | *Atilax paludinosus* | Omnivore ** | 0.97 |
| mongoose | Slender mongoose | *Herpestes sanguineus* | Omnivore ** | 0.97 |
| mongoose | White-tailed mongoose | *Ichneumia albicauda* | Omnivore ** | 0.97 |
| oribi | Oribi | *Ourebia ourebi* | Other_herbivore | 1.00 |
| porcupine | Porcupine | *Hystrix africaeaustralis* | Other_herbivore * | 1.00 |
| reedbuck | Reedbuck | *Redunca redunca* | Grazer | NA |
| serval | Serval | *Leptailurus serval* | Carnivore | 0.75 |
| hyena_aardwolf | Spotted hyaena | *Crocuta crocuta* | Carnivore | 0.95 |
| hyena_aardwolf | Striped hyaena | *Hyaena hyaena* | Omnivore | 0.95 |
| gazelle_thomsons | Thomsons gazelle | *Eudorcas thomsonii* | Grazer | 0.98 |
| topi | Topi | *Damaliscus lunatus* | Grazer | 0.88 |
| vervet_monkey | Vervet monkey | *Chlorocebus pygerythrus* | Omnivore | 1.00 |
| warthog | Warthog | *Phacochoerus africanus* | Grazer | 0.94 |
| waterbuck | Waterbuck | *Kobus ellipsiprymnus* | Grazer | 0.00 |
| wildebeest | Wildebeest | *Connochaetes taurinus* | Grazer | 0.99 |
| zebra | Zebra | *Equus quagga* | Grazer | 0.99 |
| zorilla | Zorilla | *Ictonyx striatus* | Carnivore | 1.00 |

** Broadest category including hare and springhare*

***Broadest category including all mongoose species*

***Supplementary Table 2: Prior sensitivity analyses on multispecies occupancy model (MSOM) output.*** *To assess whether prior choice strongly influenced model results, we implemented three additional models with alternative prior distributions to that of our MSOM used for inference. The posteriors obtained in this analysis were very similar among all four models, therefore we considered prior specification to have little influence over our results.*

- *Model 1 (used for species-level inference): Mean ~ normal (0,0.001); Precision = $(sd)^{-2}$; Standard deviation ~ uniform (0,2).*
- *Model 2: Mean ~ normal (0,0.001); Precision ~ gamma (0.1,0.1)*
- *Model 3: Mean ~ uniform (-20,20); Precision ~ gamma (0.1,0.1)*
- *Model 4: Mean ~ uniform (-20,20); Precision = $(sd)^{-2}$; Standard deviation ~ uniform (0,2).*

| | Model 1 | | Model 2 | | Model 3 | | Model 4 | |
|---|---|---|---|---|---|---|---|---|
| **Community-level parameter** | **Mean** | **Standard deviation** | **Mean** | **Standard deviation** | **Mean** | **Standard deviation** | **Mean** | **Standard deviation** |
| α1: proportion of open space | -0.065 | 0.114 | -0.087 | 0.119 | -0.086 | 0.118 | -0.067 | 0.112 |
| α2: shortest distance to water | -0.083 | 0.072 | -0.075 | 0.086 | -0.075 | 0.085 | -0.082 | 0.072 |
| α3: SAVI | -0.176 | 0.144 | -0.176 | 0.143 | -0.176 | 0.142 | -0.181 | 0.143 |
| α4: shortest distance to human infrastructure | -0.134 | 0.088 | -0.15 | 0.103 | -0.152 | 0.102 | -0.128 | 0.089 |
| α5: cattle trap rate | 0.356 | 0.147 | 0.357 | 0.176 | 0.361 | 0.176 | 0.322 | 0.142 |
| α6: shoat trap rate | -0.27 | 0.179 | -0.271 | 0.199 | -0.266 | 0.198 | -0.275 | 0.178 |
| α7: quadratic cattle trap rate | -0.09 | 0.043 | -0.04 | 0.074 | -0.042 | 0.072 | -0.081 | 0.041 |
| α8: shoat trap rate$^2$ | 0.027 | 0.061 | 0.024 | 0.08 | 0.024 | 0.079 | 0.029 | 0.06 |
| β1i: average grass height | -0.229 | 0.074 | -0.233 | 0.074 | -0.234 | 0.075 | -0.231 | 0.073 |
| β2i: tree/shrub density | -0.014 | 0.06 | -0.014 | 0.062 | -0.014 | 0.061 | -0.015 | 0.06 |

***Supplementary Table 3: Community-level trophic effects on occupancy (α) and detection (β) of mammal species in the Maasai Mara, Kenya.*** *Mean, standard deviation and 95% and 68% Bayesian credible intervals for two hyperparameters estimated across 20 herbivore and 15 non-herbivore species. Hyperparameters with estimates whose 95% and 68% Credible Intervals (CrI) do not overlap zero are highlighted in* ***bold*** *and marked with ** and * respectively.*

| Community-level hyper-parameter | Mean | SD | 95% Bayesian CrI | | 68% Bayesian CrI | | f |
|---|---|---|---|---|---|---|---|
| Herbivore | | | | | | | |
| **α1: proportion of open space*** | **-0.132** | 0.136 | -0.404 | 0.133 | **-0.262** | **-0.002** | 0.843 |
| α2:shortest distance to water | 0.000 | 0.090 | -0.174 | 0.177 | -0.089 | 0.088 | 0.502 |
| α3: SAVI | -0.008 | 0.216 | -0.429 | 0.428 | -0.217 | 0.201 | 0.519 |
| **α4: shortest distance to human infrastructure*** | **-0.220** | 0.137 | -0.515 | 0.032 | **-0.347** | **-0.095** | 0.959 |
| **α5: cattle trap rate*** | **0.347** | 0.204 | -0.050 | 0.745 | **0.146** | **0.548** | 0.956 |
| **α6: shoat trap rate*** | **-0.297** | 0.251 | -0.788 | 0.202 | **-0.543** | **-0.054** | 0.889 |
| **α7: quadratic cattle trap rate*** | **-0.065** | 0.066 | -0.182 | 0.074 | **-0.130** | **-0.002** | 0.847 |
| α8: shoat trap rate$^2$ | -0.012 | 0.085 | -0.188 | 0.144 | -0.093 | 0.070 | 0.547 |
| **β1i: average grass height *** | **-0.160** | 0.093 | -0.349 | 0.020 | **-0.249** | **-0.072** | 0.961 |
| β2i: tree/shrub density | -0.032 | 0.092 | -0.214 | 0.150 | -0.121 | 0.057 | 0.642 |
| Non-herbivore | | | | | | | |
| α1: proportion of open space | -0.037 | 0.289 | -0.642 | 0.521 | -0.302 | 0.229 | 0.543 |
| **α2:shortest distance to water*** | **-0.237** | 0.171 | -0.595 | 0.093 | **-0.390** | **-0.082** | 0.932 |
| **α3: SAVI**** | **-0.364** | 0.156 | **-0.685** | **-0.072** | **-0.510** | **-0.215** | 0.992 |
| α4: shortest distance to human infrastructure | -0.005 | 0.185 | -0.339 | 0.387 | -0.179 | 0.164 | 0.531 |
| **α5: cattle trap rate*** | **0.404** | 0.318 | -0.184 | 1.079 | **0.109** | **0.696** | 0.915 |
| α6: shoat trap rate | -0.286 | 0.343 | -0.990 | 0.379 | -0.616 | 0.043 | 0.805 |
| α7: quadratic cattle trap rate | -0.053 | 0.176 | -0.275 | 0.395 | -0.177 | 0.045 | 0.763 |
| α8: shoat trap rate$^2$ | 0.121 | 0.149 | -0.135 | 0.455 | -0.012 | 0.250 | 0.813 |
| **β1i: average grass height**** | **-0.387** | 0.138 | **-0.684** | **-0.131** | **-0.515** | **-0.258** | 0.997 |
| β2i: tree/shrub density | 0.003 | 0.062 | -0.118 | 0.127 | -0.057 | 0.063 | 0.522 |

***Supplementary Table 4: Species-level responses of mammals to anthropogenic pressures in Maasai Mara, Kenya, under trophic-level community effects in the Maasai Mara, Kenya.*** *Mean, standard deviation, and 95% and 68% Bayesian credible intervals for the estimates of individual species responses to three anthropogenic pressure variables in a model with distinct community-level hyperparameters based on trophic level (20 herbivore species (underlined) and 15 non-herbivore species). Estimates with 68% Credible Intervals (CrI) that do not include zero are in* ***bold*** *and denoted with* *******.

| | Human infrastructure | | | | | | Cattle | | | | | | Quadratic cattle | | | | | | Shoats | | | | | |
|---|---|---|---|---|---|---|---|---|---|---|---|---|---|---|---|---|---|---|---|---|---|---|---|---|
| **Species label** | **Mean** | **SD** | **95% CrI** | | **68% CrI** | | **Mean** | **SD** | **95% CrI** | | **68% CrI** | | **Mean** | **SD** | **95% CrI** | | **68% CrI** | | **Mean** | **SD** | **95% CrI** | | **68% CrI** | |
| aardvark | -0.050 | 0.324 | -0.704 | 0.634 | -0.310 | 0.202 | 0.269 | 0.505 | -0.779 | 1.262 | -0.149 | 0.674 | **-0.191*** | 0.301 | -0.942 | 0.177 | **-0.330** | **-0.019** | -0.206 | 0.454 | -1.074 | 0.742 | -0.608 | 0.185 |
| aardwolf | -0.034 | 0.354 | -0.748 | 0.721 | -0.306 | 0.239 | **0.736*** | 0.652 | -0.139 | 2.416 | **0.209** | **1.260** | -0.009 | 0.309 | -0.350 | 0.761 | -0.193 | 0.133 | -0.308 | 0.474 | -1.299 | 0.587 | -0.725 | 0.105 |
| baboon | 0.040 | 0.251 | -0.413 | 0.604 | -0.191 | 0.272 | 0.261 | 0.398 | -0.584 | 1.022 | -0.097 | 0.623 | -0.080 | 0.154 | -0.318 | 0.292 | -0.202 | 0.030 | -0.203 | 0.396 | -0.956 | 0.618 | -0.576 | 0.162 |
| bateared_fox | 0.004 | 0.306 | -0.565 | 0.692 | -0.250 | 0.255 | 0.326 | 0.438 | -0.524 | 1.234 | -0.055 | 0.686 | **-0.130*** | 0.223 | -0.553 | 0.246 | **-0.272** | **-0.006** | -0.272 | 0.465 | -1.235 | 0.583 | -0.674 | 0.131 |
| buffalo | **-0.501**** | 0.310 | **-1.213** | **-0.029** | **-0.808** | **-0.199** | 0.227 | 0.273 | -0.379 | 0.701 | -0.028 | 0.484 | -0.063 | 0.074 | -0.195 | 0.097 | -0.133 | 0.007 | -0.458 | 0.340 | -1.152 | 0.188 | -0.793 | -0.128 |
| bushbuck | -0.414 | 0.419 | -1.442 | 0.244 | **-0.783** | **-0.074** | **0.337*** | 0.318 | -0.299 | 0.980 | **0.062** | **0.605** | -0.050 | 0.096 | -0.204 | 0.172 | -0.134 | 0.028 | **-0.616*** | 0.515 | -1.801 | 0.237 | **-1.096** | **-0.151** |
| cheetah | 0.010 | 0.341 | -0.630 | 0.783 | -0.262 | 0.275 | 0.293 | 0.524 | -0.827 | 1.323 | -0.128 | 0.708 | -0.130 | 0.315 | -0.746 | 0.327 | -0.275 | 0.011 | -0.254 | 0.482 | -1.210 | 0.698 | -0.665 | 0.152 |
| dikdik | -0.109 | 0.284 | -0.661 | 0.501 | -0.363 | 0.153 | **0.342*** | 0.267 | -0.181 | 0.890 | **0.095** | **0.584** | -0.075 | 0.084 | -0.231 | 0.106 | -0.151 | 0.001 | -0.133 | 0.353 | -0.783 | 0.622 | -0.471 | 0.207 |
| eland | **-0.398*** | 0.305 | -1.077 | 0.124 | **-0.693** | **-0.124** | **0.463*** | 0.352 | -0.085 | 1.330 | **0.164** | **0.743** | -0.075 | 0.113 | -0.284 | 0.173 | -0.166 | 0.013 | -0.312 | 0.401 | -1.099 | 0.505 | -0.684 | 0.050 |
| elephant | 0.145 | 0.269 | -0.273 | 0.766 | -0.119 | 0.405 | **0.285*** | 0.264 | -0.267 | 0.782 | **0.037** | **0.532** | **-0.090*** | 0.083 | -0.252 | 0.074 | **-0.165** | **-0.014** | 0.121 | 0.433 | -0.588 | 1.100 | -0.292 | 0.524 |
| gazelle_grants | -0.184 | 0.240 | -0.657 | 0.327 | -0.403 | 0.032 | **0.367*** | 0.281 | -0.150 | 0.967 | **0.112** | **0.615** | **-0.081*** | 0.088 | -0.243 | 0.100 | **-0.157** | **-0.005** | **-0.496*** | 0.350 | -1.198 | 0.167 | **-0.840** | **-0.159** |
| gazelle_thomsons | **-0.358*** | 0.243 | -0.867 | 0.095 | **-0.595** | **-0.133** | **0.441*** | 0.304 | -0.084 | 1.136 | **0.169** | **0.710** | -0.033 | 0.103 | -0.185 | 0.218 | -0.122 | 0.048 | -0.214 | 0.358 | -0.889 | 0.535 | -0.552 | 0.123 |
| genet | -0.059 | 0.340 | -0.770 | 0.640 | -0.326 | 0.206 | **0.494*** | 0.521 | -0.368 | 1.740 | **0.065** | **0.916** | -0.021 | 0.299 | -0.352 | 0.716 | -0.197 | 0.111 | **-0.508*** | 0.575 | -1.926 | 0.385 | **-0.982** | **-0.020** |
| giraffe | **-0.193*** | 0.193 | -0.576 | 0.205 | **-0.375** | **-0.013** | **0.431*** | 0.293 | -0.075 | 1.100 | **0.164** | **0.687** | -0.037 | 0.099 | -0.185 | 0.196 | -0.123 | 0.044 | -0.069 | 0.356 | -0.714 | 0.690 | -0.410 | 0.275 |
| hare | 0.142 | 0.344 | -0.363 | 0.966 | -0.173 | 0.474 | **0.313*** | 0.279 | -0.243 | 0.866 | **0.056** | **0.561** | **-0.086*** | 0.086 | -0.249 | 0.085 | **-0.162** | **-0.012** | -0.048 | 0.395 | -0.737 | 0.807 | -0.417 | 0.320 |
| hartebeest_cokes | -0.098 | 0.233 | -0.530 | 0.417 | -0.307 | 0.113 | **0.301*** | 0.312 | -0.343 | 0.905 | **0.029** | **0.570** | -0.063 | 0.094 | -0.221 | 0.146 | -0.144 | 0.014 | **-0.861*** | 0.586 | -2.208 | 0.017 | **-1.398** | **-0.310** |
| hippopotamus | **-0.375*** | 0.310 | -1.112 | 0.121 | **-0.666** | **-0.101** | 0.200 | 0.306 | -0.508 | 0.705 | -0.078 | 0.480 | **-0.104*** | 0.094 | -0.320 | 0.058 | **-0.181** | **-0.022** | -0.285 | 0.358 | -1.009 | 0.423 | -0.622 | 0.054 |
| hyena_spotted | 0.105 | 0.278 | -0.309 | 0.781 | -0.137 | 0.341 | **0.556*** | 0.497 | -0.230 | 1.763 | **0.136** | **0.971** | 0.011 | 0.304 | -0.277 | 0.836 | -0.169 | 0.139 | -0.263 | 0.446 | -1.187 | 0.592 | -0.660 | 0.137 |
| hyena_striped | 0.171 | 0.408 | -0.395 | 1.198 | -0.159 | 0.509 | 0.245 | 0.591 | -1.097 | 1.324 | -0.197 | 0.699 | -0.083 | 0.312 | -0.586 | 0.552 | -0.242 | 0.053 | -0.358 | 0.551 | -1.590 | 0.611 | -0.793 | 0.090 |
| impala | -0.214 | 0.331 | -0.813 | 0.541 | -0.519 | 0.060 | **0.465*** | 0.336 | -0.071 | 1.285 | **0.175** | **0.738** | -0.045 | 0.104 | -0.204 | 0.205 | -0.132 | 0.034 | 0.112 | 0.494 | -0.675 | 1.248 | -0.342 | 0.573 |
| jackal | -0.022 | 0.264 | -0.517 | 0.552 | -0.256 | 0.207 | **0.594*** | 0.552 | -0.193 | 2.021 | **0.144** | **1.024** | -0.046 | 0.241 | -0.320 | 0.545 | -0.194 | 0.062 | -0.376 | 0.463 | -1.410 | 0.444 | -0.789 | 0.042 |
| leopard | -0.005 | 0.377 | -0.738 | 0.819 | -0.291 | 0.276 | **0.484*** | 0.542 | -0.450 | 1.753 | **0.046** | **0.914** | -0.021 | 0.309 | -0.382 | 0.789 | -0.203 | 0.113 | -0.322 | 0.536 | -1.485 | 0.662 | -0.764 | 0.120 |
| lion | -0.047 | 0.274 | -0.596 | 0.518 | -0.283 | 0.185 | 0.223 | 0.448 | -0.738 | 1.076 | -0.168 | 0.610 | -0.053 | 0.375 | -0.457 | 0.907 | -0.238 | 0.045 | **-0.483*** | 0.500 | -1.648 | 0.357 | **-0.923** | **-0.034** |
| mongoose | -0.248 | 0.290 | -0.931 | 0.214 | -0.520 | 0.020 | **0.447*** | 0.403 | -0.268 | 1.325 | **0.085** | **0.804** | -0.007 | 0.284 | -0.286 | 0.775 | -0.179 | 0.114 | -0.268 | 0.430 | -1.200 | 0.527 | -0.654 | 0.126 |
| oribi | **-0.421*** | 0.391 | -1.378 | 0.165 | **-0.774** | **-0.092** | **0.297*** | 0.285 | -0.298 | 0.835 | **0.040** | **0.556** | **-0.082*** | 0.095 | -0.275 | 0.100 | **-0.161** | **-0.001** | -0.346 | 0.422 | -1.225 | 0.471 | -0.737 | 0.041 |

| | | | | | | | | | | | | | | | | | | | | | | | | |
|---|---|---|---|---|---|---|---|---|---|---|---|---|---|---|---|---|---|---|---|---|---|---|---|---|
| porcupine | -0.281 | 0.369 | -1.117 | 0.390 | -0.600 | 0.021 | 0.238 | 0.337 | -0.546 | 0.812 | -0.043 | 0.528 | **-0.088*** | 0.103 | -0.305 | 0.106 | **-0.169** | **-0.003** | **-0.450*** | 0.461 | -1.462 | 0.392 | **-0.870** | **-0.040** |
| reedbuck | **-0.331*** | 0.392 | -1.259 | 0.330 | **-0.671** | **-0.010** | **0.312*** | 0.310 | -0.353 | 0.904 | **0.044** | **0.583** | **-0.086*** | 0.101 | -0.301 | 0.101 | **-0.166** | **-0.001** | -0.373 | 0.471 | -1.411 | 0.513 | -0.793 | 0.047 |
| serval | -0.022 | 0.328 | -0.650 | 0.709 | -0.291 | 0.235 | 0.256 | 0.477 | -0.776 | 1.159 | -0.147 | 0.657 | -0.023 | 0.298 | -0.327 | 0.691 | -0.193 | 0.092 | -0.234 | 0.466 | -1.157 | 0.728 | -0.645 | 0.173 |
| topi | -0.143 | 0.181 | -0.496 | 0.229 | -0.311 | 0.026 | **0.501*** | 0.304 | -0.005 | 1.207 | **0.222** | **0.781** | -0.020 | 0.118 | -0.181 | 0.278 | -0.117 | 0.068 | **-0.511*** | 0.331 | -1.212 | 0.086 | **-0.828** | **-0.193** |
| vervet_monkey | -0.047 | 0.231 | -0.512 | 0.427 | -0.257 | 0.160 | 0.243 | 0.359 | -0.532 | 0.918 | -0.091 | 0.580 | -0.079 | 0.114 | -0.283 | 0.163 | -0.182 | 0.021 | -0.084 | 0.423 | -0.827 | 0.869 | -0.481 | 0.303 |
| warthog | -0.104 | 0.248 | -0.532 | 0.474 | -0.324 | 0.117 | **0.392*** | 0.296 | -0.135 | 1.044 | **0.127** | **0.647** | -0.045 | 0.110 | -0.209 | 0.216 | -0.136 | 0.035 | 0.003 | 0.435 | -0.749 | 0.965 | -0.401 | 0.412 |
| waterbuck | **-0.546*** | 0.441 | -1.624 | 0.053 | **-0.957** | **-0.158** | **0.300*** | 0.290 | -0.311 | 0.848 | **0.038** | **0.559** | **-0.093*** | 0.097 | -0.301 | 0.088 | **-0.172** | **-0.010** | **-0.469*** | 0.428 | -1.404 | 0.309 | **-0.870** | **-0.082** |
| wildebeest | -0.041 | 0.328 | -0.565 | 0.743 | -0.321 | 0.245 | **0.389*** | 0.317 | -0.187 | 1.093 | **0.113** | **0.650** | -0.047 | 0.106 | -0.210 | 0.201 | -0.134 | 0.034 | -0.114 | 0.459 | -0.913 | 0.930 | -0.524 | 0.297 |
| zebra | 0.023 | 0.262 | -0.383 | 0.653 | -0.215 | 0.267 | **0.360*** | 0.283 | -0.199 | 0.944 | **0.105** | **0.616** | -0.040 | 0.097 | -0.189 | 0.191 | -0.124 | 0.038 | **-0.410*** | 0.354 | -1.099 | 0.305 | **-0.758** | **-0.075** |
| zorilla | 0.134 | 0.376 | -0.440 | 1.042 | -0.172 | 0.446 | **0.633*** | 0.534 | -0.212 | 1.880 | **0.180** | **1.094** | 0.056 | 0.307 | -0.246 | 0.863 | -0.144 | 0.225 | -0.158 | 0.485 | -1.045 | 0.862 | -0.580 | 0.253 |

### Code and data availability

On publication, all code will be available on GitHub. For peer review, we provide an anonymised version of code for the Maasai Mara Classifier (MMC) model here: https://anonymous.4open.science/r/camera_traps_classifier-7F51. R code to run our multispecies occupancy model along with detection matrices and covariates are also available on this repository. On publication, training, test, and validation images will be available on LILA BC (lila.science).
